# foxQ2 evolved a key role in anterior head and central brain patterning in protostomes

**DOI:** 10.1101/090340

**Authors:** Peter Kitzmann, Matthias Weibkopf, Magdalena Ines Schacht, Gregor Bucher

## Abstract

Anterior patterning of animals is based on a set of highly conserved transcription factors but the interactions within the protostome anterior gene regulatory network (aGRN) remain enigmatic. Here, we identify the *foxQ2* ortholog of the red flour beetle *Tribolium castaneum* as novel upstream component of the insect aGRN. It is required for the development of the labrum and higher order brain structures, namely the central complex and the mushroom bodies. We reveal *Tc-foxQ2* interactions by RNAi and heat shock-mediated misexpression. Surprisingly, *Tc-foxQ2* and *Tc-six3* mutually activate each other forming a novel regulatory module at the top of the insect aGRN. Comparisons of our results with those of sea urchins and cnidarians suggest that *foxQ2* has acquired functions in head and brain patterning during protostome evolution. Our findings expand the knowledge on *foxQ2* gene function to include essential roles in epidermal development and central brain patterning.

**Author summary:** The development of the anterior most part of any animal embryo – for instance the brain of vertebrates and the head of insects – depends on a very similar set of genes present in all animals. This is true for the two major lineages of bilaterian animals, the deuterostomes (including sea urchin and humans) and protostomes (including annelids and insects) and the cnidarians (e.g. the sea anemone), which are representatives of more ancient animals. However, the interaction of these genes has been studied in deuterostomes and cnidarians but not in protostomes. Here, we present the first study the function of the gene *foxQ2* in protostomes. We found that the gene acts at the top level of the genetic network and when its function is knocked down, the labrum (a part of the head) and higher order brain centers do not develop. This is in contrast to the other animal groups where *foxQ2* appears to play a less central role. We conclude that *foxQ2* has acquired additional functions in the course of evolution of protostomes.

## Introduction

Anterior patterning in bilaterian animals is based on a set of highly conserved transcription factors like *orthodenticle*/*otx*, *empty spiracles*/*emx*, *eyeless*/*Pax6* and other genes, which have comparable expression and function from flies to mice (Hirth et al., 1995; Leuzinger et al., 1998; Quiring et al., 1994; Simeone et al., 1992). Likewise, canonical Wnt signaling needs to be repressed in order to allow anterior pattern formation in most tested animals with *Drosophila* being a notable exception (Fu et al., 2012; Martin and Kimelman, 2009; Petersen and Reddien, 2009). Since recently, additional genes have been studied, which are expressed anterior to the *orthodenticle/otx* region at the anterior pole of embryos of all major clades of Bilateria and their sister group, the Cnidaria. These data suggested that a distinct but highly conserved anterior gene regulatory network (aGRN) governs anterior-most patterning (Lowe et al., 2003; Marlow et al., 2014; Posnien et al., 2011b; Sinigaglia et al., 2013; Steinmetz et al., 2010; Yaguchi et al., 2008). This region gives rise to the apical organ in sea urchins, annelids and hemichordates and other structures. Most of the respective orthologs were shown to be expressed in the vertebrate anterior neural plate and the anterior insect head as well (Posnien et al., 2011b) although it remains disputed what tissue-if any-is homologous to the apical organ in these species (Hunnekuhl and Akam, 2014; Marlow et al., 2014; Santagata et al., 2012; Sinigaglia et al., 2013; Telford et al., 2008). Unfortunately, detailed interactions of this aGRN were determined only in a few model systems (Range and Wei, 2016; Sinigaglia et al., 2013; Yaguchi et al., 2008).

The aGRN of sea urchins as representative of deuterostomes is best studied in *Strongylocentrotus purpuratus* and *Hemicentrotus pulcherrimus*. Data indicate that sea urchin *six3* is the most upstream regulator, which is initially co-expressed with *foxQ2*. Both genes are restricted to the anterior pole by repression by posterior Wnt signaling (Range and Wei, 2016; Wei et al., 2009; Yaguchi et al., 2008). *six3* in turn is able to repress Wnt signaling (Wei et al., 2009) and to activate a large number of genes including *rx, nk2.1* and *foxQ2* (Wei et al., 2009). Subsequently, *foxQ2* represses *six3* but activates *nk2.1* expression at the anterior-most tip. In this tissue freed of *six3* expression *foxQ2* is responsible for establishing a signaling center involved in the differentiation of the apical organ (Range and Wei, 2016). In addition, *foxQ2* expression is positively regulated by nodal signaling (Yaguchi et al., 2016). *six3* knockdown leads to a strong morphological phenotype including change of embryonic epidermal shape and loss of neural cells (Wei et al., 2009). In *foxQ2* knockdown, in contrast, an epidermal phenotype other than epidermal animal plate thickening was not observed (Yaguchi et al., 2008). In contrast, *foxQ2* appears to be essential for the specification of specific neural cell types (Yaguchi et al., 2008; Yaguchi et al., 2012; Yaguchi et al., 2016).

The *Nematostella vectensis* (Cnidaria) aGRN has been studied representing the sister group to bilaterian animals (Sinigaglia et al., 2013). Both *Nv-six3* and *Nv-foxQ2* are repressed by posterior Wnt signaling and *Nv-six3* activates *Nv-foxQ2* and a number of other genes of the aGRN (Marlow et al., 2013; Sinigaglia et al., 2013). Like in the sea urchin, knockdown of *Nv-six3* leads to strong morphological defects including loss of the apical organ while *Nv-foxQ2* knockdown does not affect the morphology of the embryo but the apical organ is reduced (Sinigaglia et al., 2013). One notable difference of the aGRN is that *Nv-foxQ2* does not appear to regulate *Nv-six3*. The repression of *six3* and *foxQ2* by Wnt signaling has been confirmed in a hemichordate (deuterostome) (Darras et al., 2011; Fritzenwanker et al., 2014) and for *foxQ2* in a hydrozoan (cnidarian) (Momose et al., 2008). Neither sea urchins nor cnidarians possess a highly developed CNS such that a function of *foxQ2* in brain development could not be tested.

Within protostomes, the expression of aGRN genes has been studied extensively in postembryonic stages of the annelid *Platynereis dumerilii* (Lophotrochozoa) but the only functional interaction tested is the repression of *Pd-six3* and *Pd-foxQ2* by Wnt signaling (Marlow et al., 2014). Within arthropods (Ecdysozoa) the red flour beetle *Tribolium castaneum* has become the main model system for studying anterior patterning (Posnien et al., 2010), because head development is more representative for insects than the involuted head of *Drosophila*. Indeed, the respective aGRNs differ significantly. The anterior morphogen *bicoid* is present only in higher dipterans while the canonical anterior repression of Wnt signaling is observed in *Tribolium* only (Brown et al., 2001; Stauber et al., 1999). Neither terminal Torso signaling nor the terminal gap gene *huckebein* have an influence on head formation in *Tribolium* (Schoppmeier and Schröder, 2005; Kittelmann et al., 2013). Apparently, there are two anterior patterning systems – one with a high degree of similarity with vertebrate neural plate patterning and another comprised by a different set of genes patterning the largely non-neural anterior median region (AMR) (Kittelmann et al., 2013; Posnien et al., 2011b). Interestingly, *Tc-six3* is a central regulator of anterior head and brain patterning, which is repressing *Tc-wg* expression (i.e. Wnt signaling) among other genes but is not regulating *Tc-rx* and *Tc-nk2.1* (Posnien et al., 2011b).

In an ongoing genome-wide RNAi screen in *Tribolium* (iBeetle screen) (Dönitz et al., 2015; Schmitt-Engel et al., 2015) a head phenotype similar to the one of *Tc-six3* was induced by the dsRNA fragment *iB_03837*. The targeted gene was the *Tribolium* ortholog of FoxQ2 (*Tc-foxQ2*), which is a Forkhead transcription factor. All members of this family share the Forkhead DNA-binding domain and they are involved in development and disease (Benayoun et al., 2011). While being highly conserved among animals, this gene was lost from placental mammals (Mazet et al., 2003; Yu et al., 2008). Within arthropods, anterior expression was described for the *Drosophila* ortholog *fd102C* (*CG11152*) (Lee and Frasch, 2004) and *Strigamia maritima* (myriapod) *foxQ2* (Hunnekuhl and Akam, 2014). However, the function of this gene has not been studied in any protostome so far.

We studied expression and function of *Tc-foxQ2* by RNAi and heat shock-mediated misexpression. We found that *Tc-foxQ2* has acquired a much more central role in the insect aGRN compared to sea urchin and cnidarians. Surprisingly, *Tc-foxQ2* and *Tc-six3* form a regulatory module with mutual activation contrasting the clear upstream role of *six3* in the other species. Another marked difference is that *Tc-foxQ2* knockdown led to a strong epidermal phenotype. Further, we found a novel role of *Tc-foxQ2* in insect central brain patterning. Specifically, it was required for the development of the mushroom bodies (MB) and the central complex (CX), both of which are higher order processing centers of the insect brain (Heisenberg, 2003; Pfeiffer and Homberg, 2014). Finally, we present the most comprehensive aGRN available for any protostome.

## Results

### *Tc-foxQ2* - a novel player in anterior head development of Tribolium

In the iBeetle screen, injection of the dsRNA fragment *iB_03837* led to first instar larval cuticles with reduced or absent labrum with high penetrance (Dönitz et al., 2015; Schmitt-Engel et al., 2015) (Fig. 1). The targeted gene was *TC004761* (Tcas_OGS 3.0), which we determined by phylogenetic analysis to be the *Tribolium* ortholog of FoxQ2 (Tc-FoxQ2) (Fig. S1). Quantitative analyses of parental RNAi experiments with two non-overlapping dsRNA fragments (*Tc-foxQ2*^RNAi_a^; *Tc-foxQ2*^RNAi_b^; 1.5 µg/µl) (Fig. 1A,B; Tables S1-4) in two different genetic backgrounds (Fig. S2; Tables S3-6) revealed the same morphological phenotype arguing against off-target effects or strong influence of the genetic background (Kitzmann et al., 2013). The proportions of eggs without cuticles or with cuticle remnants (*strong defects*) were within in the range observed in wild-type (wt). Weak *Tc-foxQ2* RNAi phenotypes showed a labrum reduced in size and loss of one or both labral setae (Fig. 1E,F; yellow dots). Intermediate phenotypes were marked by a reduced labrum and loss of one or both anterior vertex triplet setae indicating additional deletions of the head capsule (Fig. 1E,G; red dots). In strong phenotypes, the labrum was strongly reduced or deleted along with several setae marking the anterior head and/or the labrum (Fig. 1H). No other specific L1 cuticle phenotypes were detected. We tested higher dsRNA concentrations (2 µg/µl and 3.1 µg/µl; data not shown) as well as double RNAi using both dsRNA fragments together (*Tc-foxQ2*^RNAi_a^ and *Tc-foxQ2*^RNAi_b^, each 1.5 µg/µl; data not shown). None of these variations resulted in a stronger cuticle phenotype. Taken together, *Tc-foxQ2* is required for epidermal patterning of anterior head structures. Interestingly, the RNAi phenotype was very similar to the one described for *Tc-six3* although the affected domain was a bit smaller the penetrance was lower (Posnien et al., 2011b).

**Figure 1:**
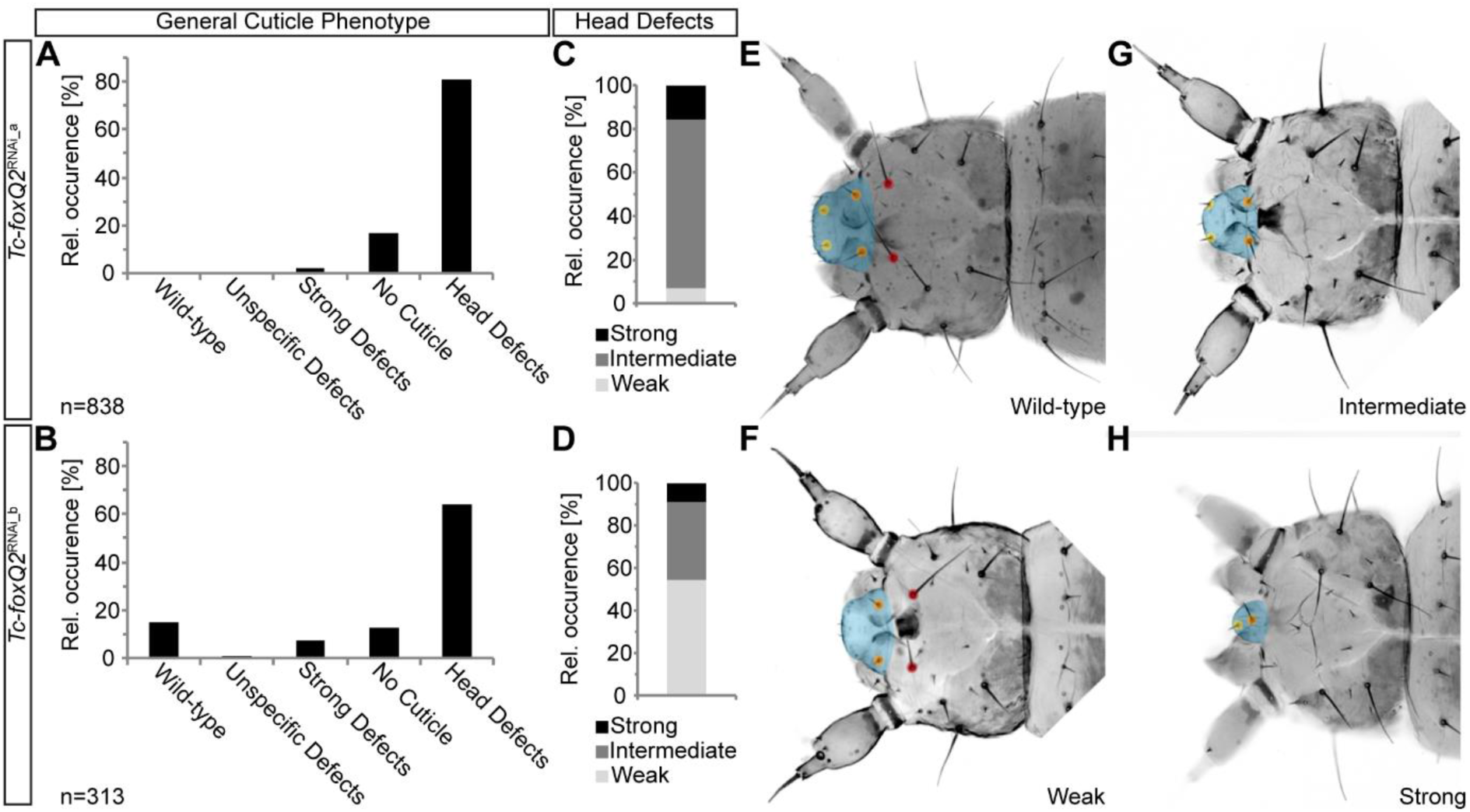
Quantitative analysis of the *Tc-foxQ2*^RNAi^ epidermal L1 defects confirms the phenotype and excludes off-target effects. (**A**, **B**) Knockdown of *Tc-foxQ2* with two non-overlapping dsRNA fragments *Tc-foxQ2*^RNAi_a^(**A**) *and Tc-foxQ2*^RNAi_b^ (**B**) leads to comparable portions of cuticle phenotypes. (**C**, **D**) Detailed analysis of the head defects shows that the *Tc-foxQ2*^RNAi_a^ dsRNA fragment leads to a qualitatively comparable but quantitatively stronger phenotype, marked by more intermediate (**G**) and strong (**H**) head defects. (**E-H**) L1 cuticle heads, depicted in a dorsal view, are representing the different classes of head defects. Anterior is left. (**E**) Wt cuticle with the labrum marked in blue, two labrum setae (yellow dots), two clypeus setae (orange dots) and two anterior vertex setae (red dots). (**F**) Weak head defect with a reduced labrum and at least one deleted labrum seta. (**G**) Intermediate head defect is additionally lacking at least one of the anterior vertex seta. (**H**) Strong head defect with a strongly reduced labrum, one labrum seta and one clypeus seta. The strongest phenotypes show a completely absent labrum and deleted anterior vertex setae.

### Increased apoptosis in *Tc-foxQ2* RNAi

The labrum is an appendage-like structure (Posnien et al., 2009a) and its outgrowth requires cell proliferation regulated by *Tc-serrate (Tc-ser)* (Siemanowski et al., 2015). We asked when the size of the labrum decreased after *Tc-foxQ2* RNAi and whether cell proliferation or cell death were involved. We found that the labral buds in *Tc-foxQ2*^RNAi^ embryos were decreased from fully elongated germ band stages onwards (Fig. 2Aa`,Ab`) and they fused precociously (Fig. 2Ac`,Ad`). We found no regulation of *Tc-ser* by *Tc-foxQ2* (see below). Next, we quantified apoptosis in embryos (6-26 h after egg laying (AEL); Table S7) using an antibody detecting cleaved *Drosophila* death caspase-1 (Dcp-1) (Florentin and Arama, 2012). We quantified apoptotic cells in the labral region (region 1 in Fig. 2B) and a control region (region 3 in Fig. 2E). We found that fully elongated germ bands showed a six times increased number of apoptotic cells after *Tc-foxQ2* RNAi (*p=*0.00041, n=15; Fig. 2C) coinciding with the stage of morphological reduction. Hence, *Tc-foxQ2* prevents apoptosis in the growing labrum.

**Figure 2:**
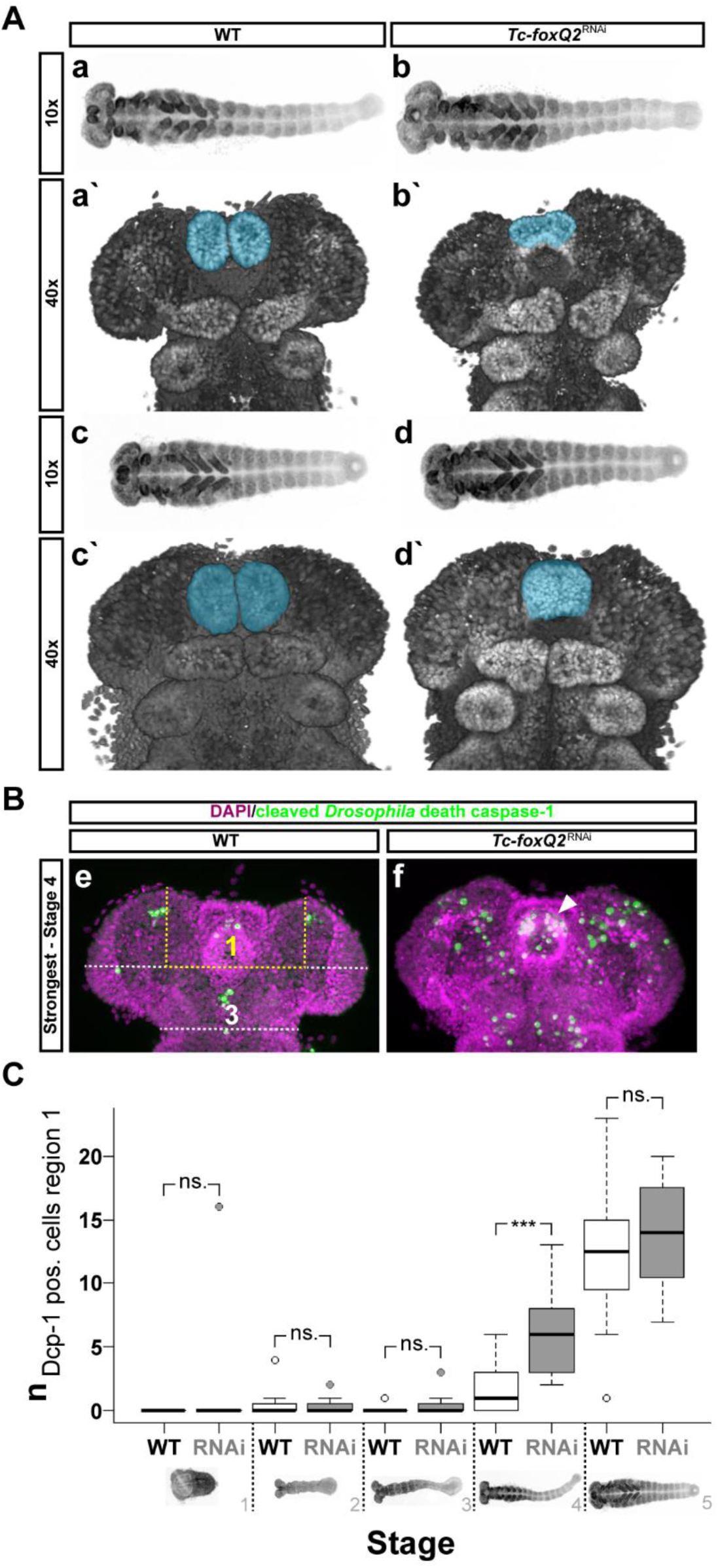
Emergence of the *Tc-foxQ2*^RNAi^ phenotype and cell death rates. (A) Morphology of wt (A_a_, A_a`_, A_c_, A_c`_) and *Tc-foxQ2*^RNAi^ (A_b_, A_b`_, A_d_, A_d`_) embryos is visualized by nuclear staining (DAPI, grey). Anterior is left in 10x panels (A_a-d_) and up in 40x panels (A_a`_). The labrum is marked in blue (A_a`-d`_). (A_a-_d) *Tc-foxQ2*^RNAi^ embryos (6-26 h AEL) show decreased labral buds, which appear to fuse prematurely(A_b`_, A_d`_). (B) For quantification of cell death rates, a region of interest (ROI) and a control region were defined. The region of interest is the labral region (region 1, yellow dashed lines). The posterior procephalic region was used for data normalization (region 3, white dashed lines). Morphology of wt (B_e_) and *Tc-foxQ2*^RNAi^ (B_f_) embryos is visualized by nuclear staining (DAPI, magenta). Apoptotic cells are monitored by antibody staining (cleaved *Drosophila* death caspase-1 (Dcp-1) - Alexa Fluor 488; green). (B_e_, B_f_) The fully elongated *Tc-foxQ2*^RNAi^ germ band with the most apoptotic cells within the labral region (ROI 1, marked with dashed lines) showed apparently more marked cells (B_f_) than the strongest representative of the wt embryos within this stage and region (B_e_). (B_f_) The *Tc-foxQ2*^RNAi^ embryo showed an accumulation of apoptotic cells within the labral buds (arrowhead). (C) Box plot depicting the normalized number of apoptotic cells (y-axis) versus five different embryonic stages, subdivided in untreated (wt) and *Tc-foxQ2*^RNAi^ embryos (x-axis). The ROI 1 values are normalized with the region 3 values (Be). Brackets display grade of significance. Germ rudiments (stage 1) to intermediate elongating germ bands (stage 3), as well as early retracting germ bands (stage 5) show no significant increase in the number of apoptotic cells within the ROI 1 of *Tc-foxQ2*^RNAi^ embryos (stage 1: *p*=0.33 (wt: n=3, RNAi: n=7), stage 2: *p*=0.63 (wt: n=11, RNAi: n=12), stage 3: *p*=0.19 (wt: n=9, RNAi: n=19), stage 5: *p*=0.15 (wt: n=12, RNAi: n=11)). However, fully elongated germ bands showed significantly more apoptotic cells (*p*=0.00041) in *Tc-foxQ2*^RNAi^ embryos (n=15) compared to untreated ones (n=17). ns.: not significant

### *Tc-foxQ2* is required for brain development

The phenotypic similarity to *Tc-six3*^RNAi^ embryos (Posnien et al., 2011b) and the expression of *Tc-foxQ2* in neuroectodermal tissue (see below) prompted us to check for brain phenotypes. We performed parental RNAi in the background of the double transgenic *brainy* line, which marks glia and neurons (visualized in white and yellow in Fig. 3A-C, respectively) (Koniszewski et al., 2016). In the weakest phenotypes the medial lobes of the MBs were reduced and appeared to be medially fused (Fig. 3, compare cyan arrowheads in A,A´ with B,B´; MBs marked in magenta). Further, the CB (part of the CX) was shortened (Fig. 3B,B´, CB marked in yellow) and the brain hemispheres were slightly fused (Fig. 3, compare black arrowheads in B´,C´ with A´). In stronger phenotypes, the CB was clearly reduced in size and the MBs were not detectable anymore. Further, the brain hemispheres appeared fused at the midline (Fig. 3C,C´). We tested the MB phenotype by RNAi in the background of the transgenic *MB-green* line which marks MBs by EGFP (Koniszewski et al., 2016). We observed a similar range of MB body phenotypes (Fig. S3). In addition, we found misarranged MBs had lost their medial contact (Fig. S3, filled arrowheads). Noteworthy, the strength of epidermal and neural phenotypes correlated. Larvae with weak neural defects showed a decreased labrum, while strong neural phenotypes correlated with lack of the entire labrum. Taken together, we found *Tc-foxQ2* to be required for brain formation with the MBs, the CB (part of the CX) and the midline being strongly affected. Again, these defects are similar to those reported for *Tc-six3* loss-of-function (Posnien et al., 2011b).

**Figure 3:**
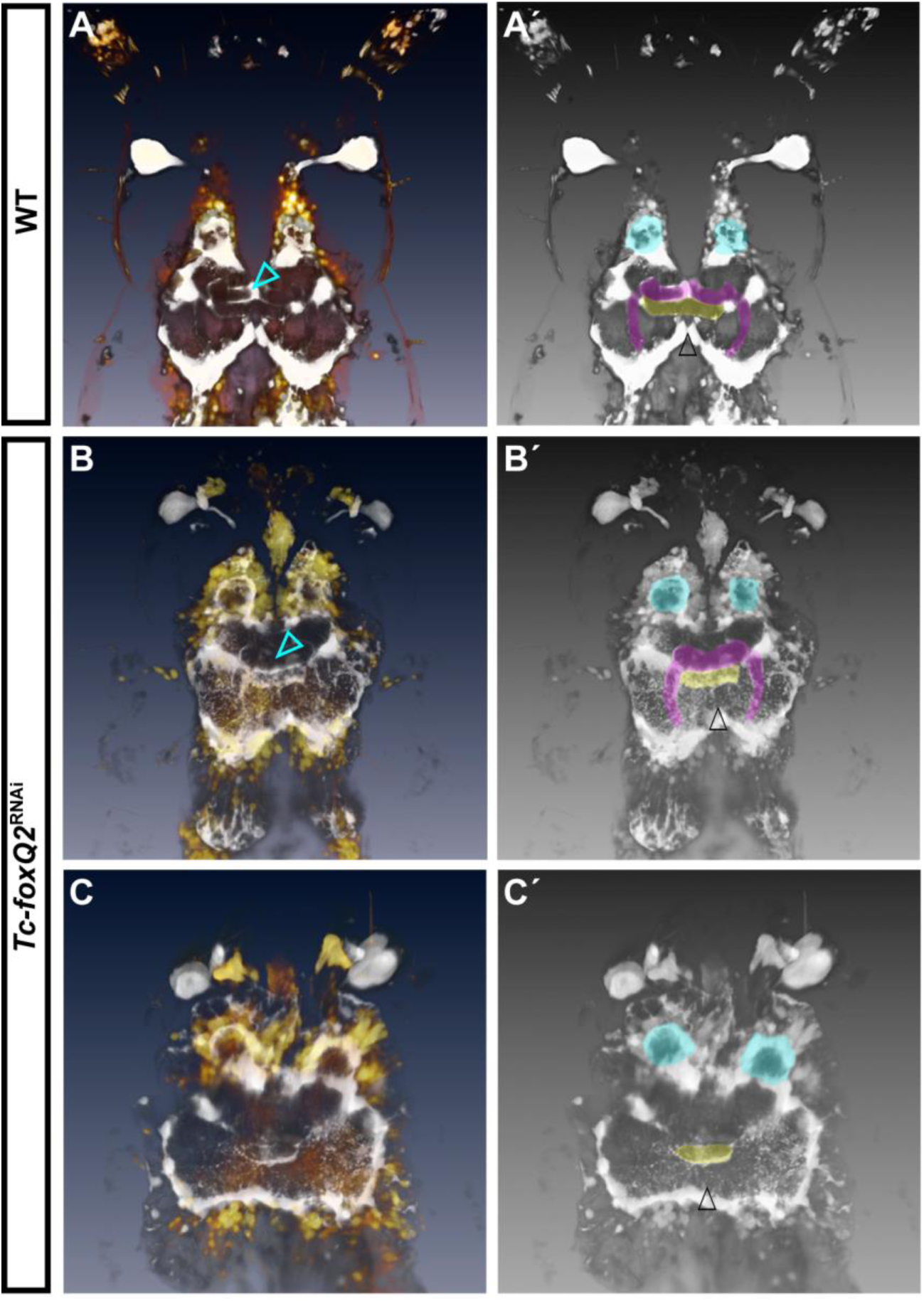
Embryonic loss of *Tc-foxQ2* function leads to defects in L1 larval brains. Anterior is up. Neural cells (yellow) and glial tissue (white) are, in wt (**A**, **A**) and *Tc-foxQ2*^RNAi^ (**B**-**D**) L1 larvae visualized by using the double transgenic *brainy* reporter line. The grey-scale images are for a better visualization of the regions of interest. (**A**, **A**) Wt L1 larval brain that shows the two brain hemispheres, each with a mushroom body (magenta), an antennal lobe (cyan), and the mid-line spanning central body (yellow). (**B**, **B**) A weak *Tc-foxQ2*^RNAi^ larval brain phenotype showing the loss of the boundary between the medial lobes of the mushroom bodies (compare cyan arrowheads in **B** and **A**). The central body appears to be slightly reduced in size. (**C**, **C**) Intermediate *Tc-foxQ2*^RNAi^ larval brains appear to lack the complete mushroom bodies. The central body appears to be reduced in size.

### Dynamic expression of *Tc-foxQ2* in the anterior head

The expression of *Tc-foxQ2* started in two domains at the anterior terminus of the germ rudiment (Fig. 4A,B). At late elongating germ band stages, the domains split into several subdomains in the AMR like in the labrum anlagen (empty arrowhead in Fig. 4F,H) and domains lateral to the stomodeum (empty arrow in Fig. 4G,I). Besides, there are also domains in the neuroectoderm (e.g. white arrow in Fig. 4F,L; see approximate fate map in Fig. S4). Very weak staining in the ocular region was detected with the tyramide signal amplification (TSA) system (Fig. 4K). These data confirm the anterior expression of *foxQ2* orthologs found in other animals and is in line with its function in labrum and neural development.

**Figure 4:**
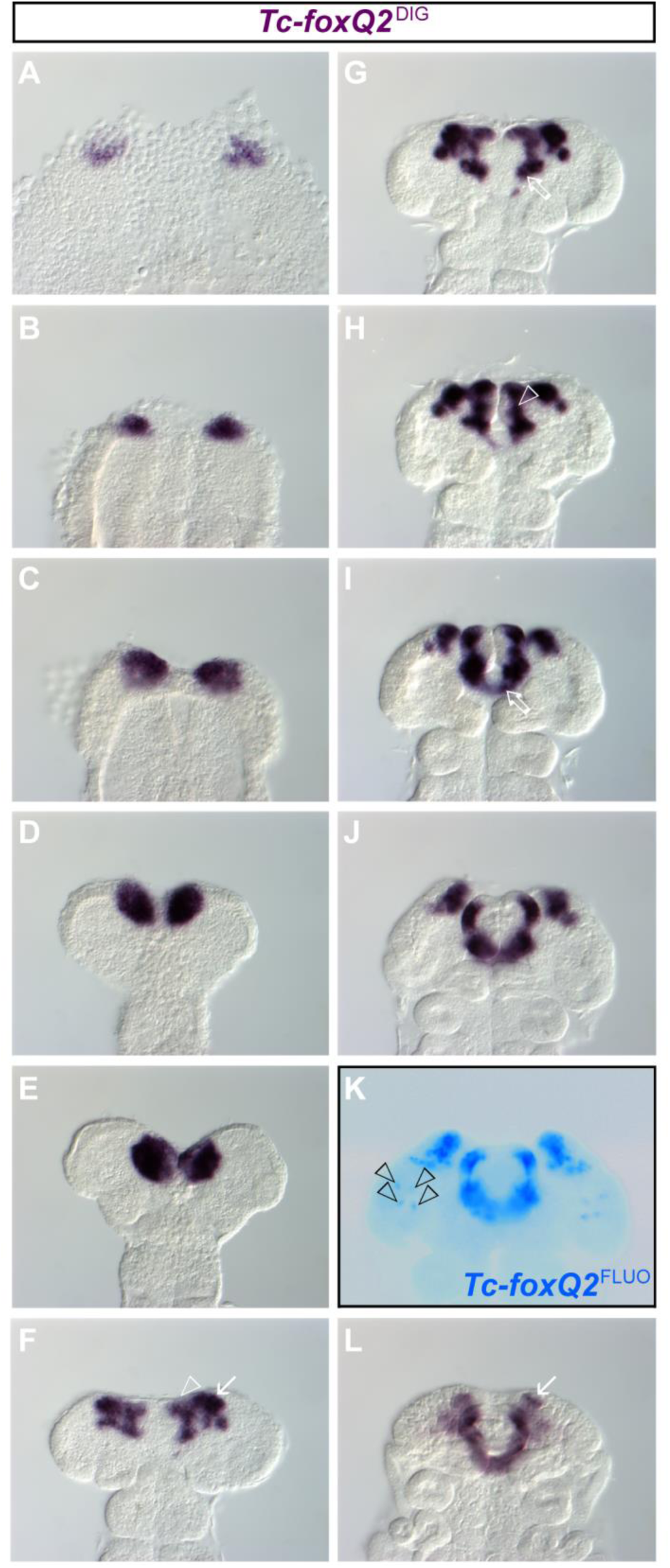
*Tc-foxQ2* is expressed in a highly dynamic pattern at the anterior pole. Anterior is up. Expression of *Tc-foxQ2* in wild-type (wt) embryos is monitored by whole mount in situ hybridization (ISH). (**A**) *Tc-foxQ2* expression starts with the formation of the germ rudiment. (**A**-**E**) The early *Tc-foxQ2* expression is marked by two domains located at the anterior pole, which successively approach each other, probably due to morphogenetic movements, at the embryonic midline. (**F**) The expression pattern splits into several domains in late elongating germ bands, with expression domains in the putative neuroectoderm (white arrow, presumably including parts of the pars intercerebralis; Posnien 2011) and in the labral/stomodeal region (white empty arrowhead). (**G**) The expression domains flanking the prospective stomodeum become more prominent (empty arrow). (**H**) The anterior median expression domain frames the lateral parts of the labral buds (white empty arrowhead). (**I**, **J**) At fully elongated to early retracting germ band stages the two expression domains flanking the stomodeum are posteriorly linked to each other (empty arrow) and the expression domains within the labral buds are getting narrower. (**K**) The staining with the more sensitive TSA-Dylight550 (*Tc-foxQ2*^FLUO^) reveals four dot-like expression domains in the ocular region (black empty arrowheads). (**L**) At retracting germ band stages *Tc-foxQ2* is expressed in a narrow U-shape pattern and the neuroectodermal expression domains are reduced in size (white arrow).

We mapped the expression of *Tc-foxQ2* relative to other genes of the aGRN by double in situ hybridization in wt embryos (6-26 h AEL). Expression overlaps are marked with dashed lines in Figs 5 & 6. First we tested for genes that were previously described to interact with *foxQ2* in other species (see introduction) or are required for labrum formation (Coulcher and Telford, 2012; Economou and Telford, 2009; Kittelmann et al., 2013; Posnien et al., 2009a; Posnien et al., 2011b). We found no overlap with *Tc-wingless/wnt1* (*Tc-wg*) expression until retraction, where the emerging stomodeal *Tc-wg* domain overlapped with *Tc-foxQ2* expression (Fig. 5A). In contrast, we found complete overlap with *Tc-six3* expression at early embryonic stages (Fig.5B_0_), which developed into a mutually exclusive expression at intermediate elongating germ bands (Fig. 5B_2_). Afterwards, these genes remained expressed mutually exclusive apart from a small anterior median neuroectodermal region (lateral area marked in Fig. 5B_3-6_) and the labrum anlagen (median area marked in Fig. 5B_5-6_). These data are in agreement with the interactions described for sea urchin where *six3* initially activates *foxQ2* while at later stages *six3* gets repressed by *foxQ2* (see introduction). The later coexistence of mutually exclusive and co-expression domains along with the many different expression domains of *Tc-foxQ2* indicate a complex and region specific regulation. Early co-expression developing into partially overlapping expression patterns was observed for both *Tc-cap´n´collar* (*Tc-cnc*) and *Tc-scarecrow* (*Tc-scro*/*nk2.1*) (Fig. 5C-D). For *Tc-crocodile* (*Tc-croc*) a small overlap was observed, which remained throughout development (Fig. 5E). (Coulcher and Telford, 2012; Economou and Telford, 2009; Kittelmann et al., 2013; Posnien et al., 2011b).

**Figure 5:**
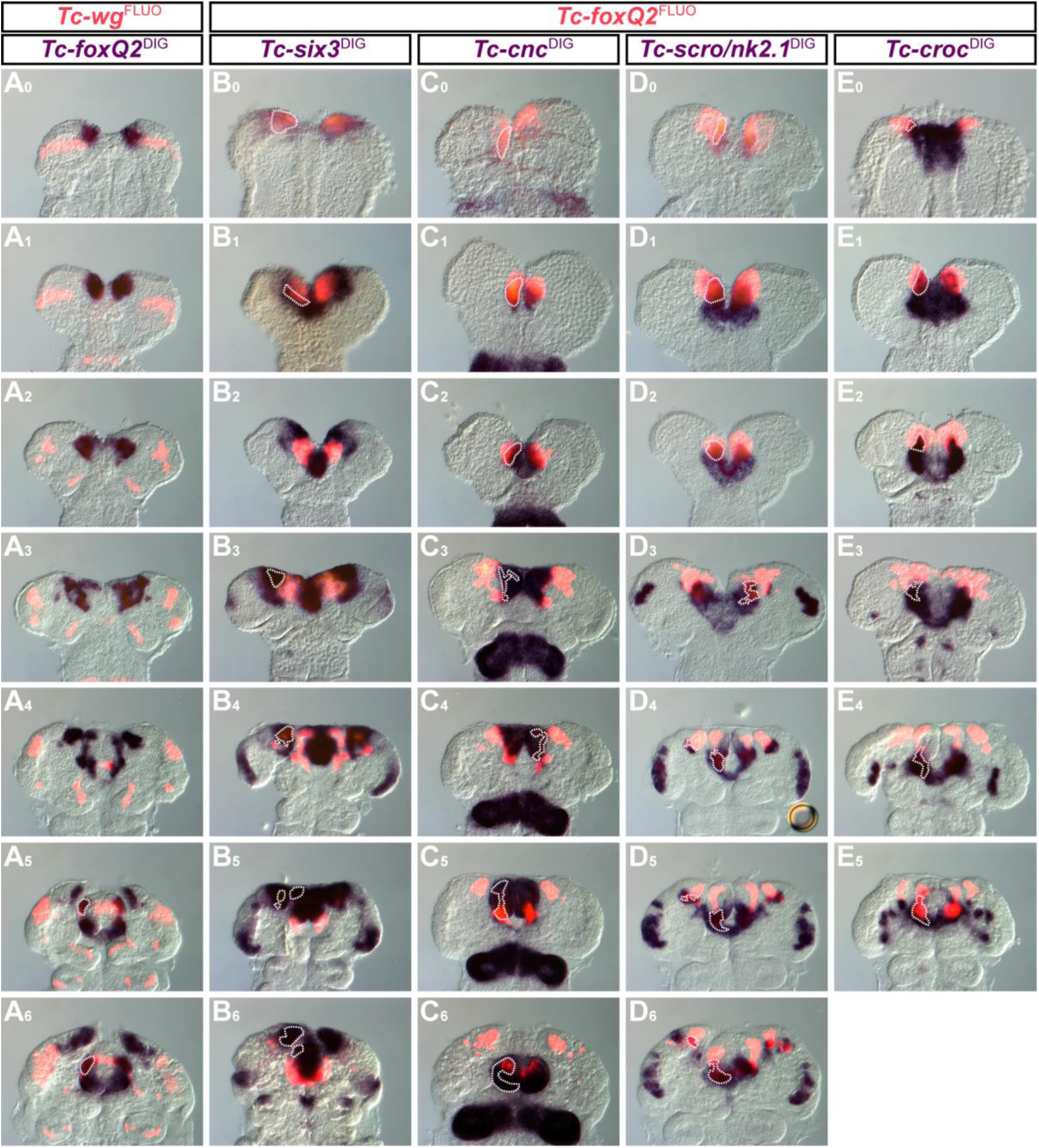
Co-expression analyses of *Tc-foxQ2* and anterior head patterning genes I. Anterior is up. Expression is visualized by double ISH (DISH), using NBT/BCIP (blue) and TSA-Dylight550 (red). For better comparison and potential TSA signal quenching effects by NBT/BCIP staining, *Tc-foxQ2* was stained using NBT/BCIP in **A_0_**_-**6**_. Co-expression is indicated with dashed lines. (**A_0_**_-**6**_) Until fully elongated germ band stages no *Tc-foxQ2*/*Tc-wg* co-expression is detectable (**A_0_**_-**4**_). At later stages *Tc-foxQ2* is co-expressed with *Tc-wg* in the anterior portion of the labral buds (**A_5_**, **A_6_**). (**B_0_**_-**6**_) *Tc-foxQ2* and *Tc-six3* are completely overlapping in their expression at germ rudiment stages (**B_0_**). In early elongating germ bands the co-expression is limited to a narrow lateral stripe of the AMR (**B_1_**). Intermediate germ bands show a mutually exclusive expression of *Tc-foxQ2* and *Tc-six3*, within the AMR (**B_2_**). At later stages *Tc-foxQ2* and *Tc-six3* expression are overlapping within the neurogenic region (**B_3_**_-**4**_). In early retracting germ bands and at later stages *Tc-foxQ2* and *Tc-six3* expression additionally overlap in the anterior portion of the labral buds (**B_5_**_-**6**_). (**C_0_**_-**6**_) *Tc-cnc* expression is almost completely covering the expression of *Tc-foxQ2*, at early embryonic stages (**C_0_**_-**2**_). At later stages the co-expression is restricted to parts of the labral/stomodeal region (**C_3_**_-**6**_). (**D_0_**_-**6**_) *Tc-scro*/*nk2.1* is partially co-expressed with *Tc-foxQ2* within the anterior part of the AMR at early embryonic stages (**D_0_**_-**2**_). In late elongating germ bands the co-expression is restricted to a narrow lateral stripe of the AMR (**D_3_**). Later stages show co-expression of *Tc-scro*/*nk2.1* and *Tc-foxQ2* in the posterior portion of the labral buds, the stomodeum flanking region and small areas of the neurogenic region (**D_4_**_-**6**_). *Tc-croc* expression is partially overlapping with *Tc-foxQ2* within antero-lateral parts of the AMR at early stages (**E_0_**_-**2**_). At later stages of development the *Tc-foxQ2*/*Tc-croc* co-expression is restricted to the stomodeum flanking region and the posterior portion of the labral buds (**E_3_**_-**5**_).

*Tc-retinal homeobox* (*Tc-rx*) was expressed in a largely non-overlapping pattern apart from small domains in labrum and neuroectoderm at late stages (Fig. 6A). *Tc-chx* and *Tc-ser* start expression largely overlapping with *Tc-foxQ2* but later resolve to mainly non-overlapping patterns (Fig. 6B,E). *Tc-forkhead (Tc-fkh/foxA)* expression was essentially non-overlapping (Fig. 6C). *Tc-six4* marks a region with molecular similarity to the vertebrate placodes (Posnien et al., 2011a). No co-expression was observed until late stages where a small domain in the anterior median neuroectoderm expresses both genes (Fig. 6D). In summary, *Tc-foxQ2* expression indicated a central and dynamic role in the aGRN.

**Figure 6:**
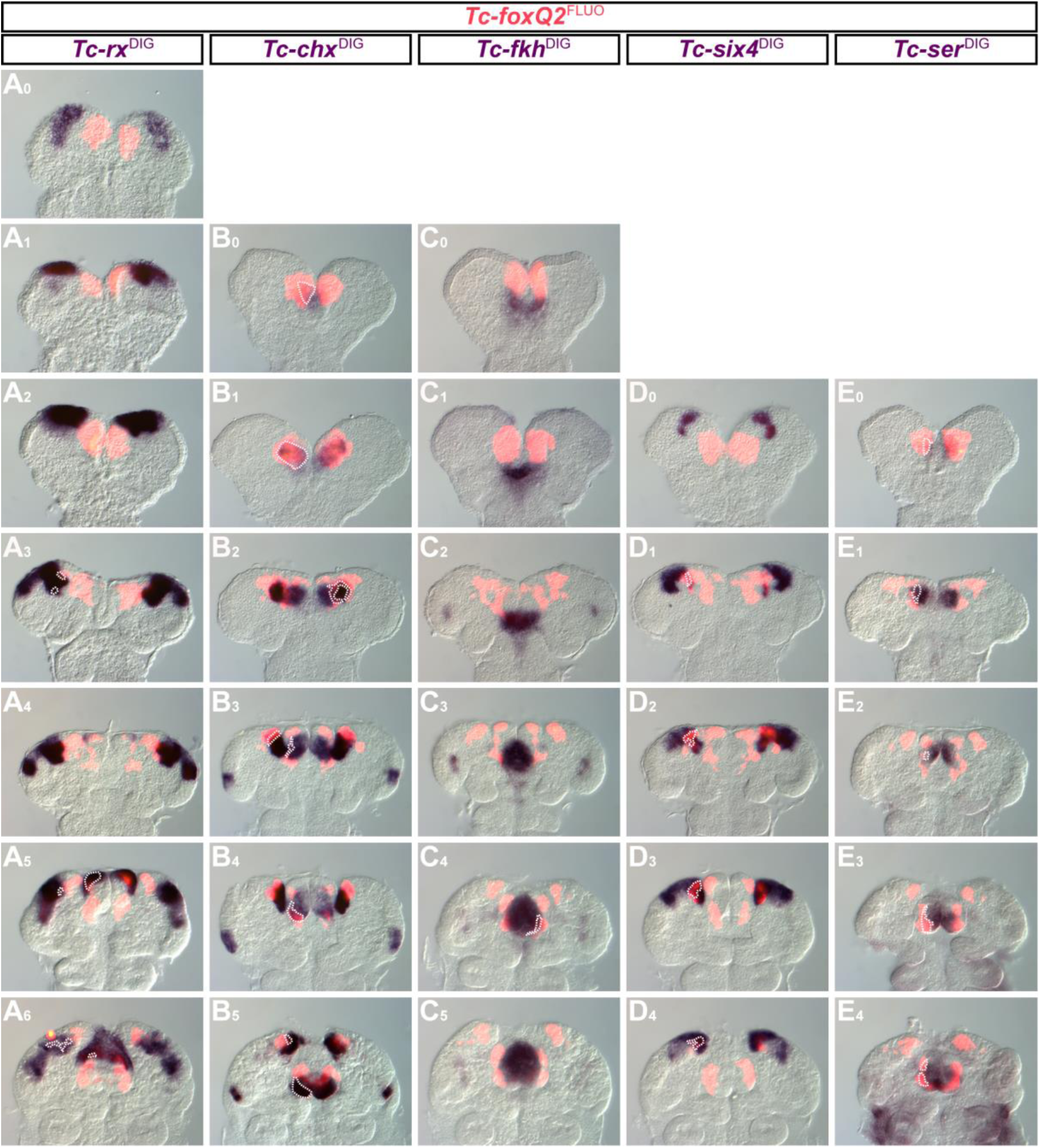
Co-expression analyses of *Tc-foxQ2* and anterior head patterning genes II. Anterior is up. Expression is visualized by DISH, using TSA-Dylight550 (red) and NBT/BCIP (blue). Co-expression is indicated with dashed lines. (**A_0_**_-**6**_) *Tc-rx* is not co-expressed with *Tc-foxQ2* until fully elongated germ band stages (**A_0_**_-**4**_), except for two little spots in the neurogenic region in late elongating germ bands (**A_3_**). In retracting germ bands both genes are partially overlapping within the neurogenic region and in anterior parts of the labral buds (**A_5_**, **A_6_**). (**B_0_**_-**5**_) *Tc-chx* expression is partially (**B_0_**) and later almost completely (**B_1_**) overlapping with *Tc-foxQ2* expression. At later stages the co-expression is restricted to narrow stripes within the outer lateral labral and the neurogenic region (**B_2_, B_3_**). In early retracting germ bands *Tc-chx* expression shows only a little overlap within the posterior portion of the labral buds (**B_4_**) and at later stages an additional overlap within the neurogenic region (**B_5_**). (**C_0_**_-**5**_) *Tc-fkh* shows almost no co-expression with *Tc-foxQ2*, except for a small domain in the stomodeal region, in early retracting germ bands (**C_5_**). (**D_0_**_-**4**_) *Tc-six4* is not co-expressed with *Tc-foxQ2*, in intermediate elongating germ bands (**D_0_**). Co-expression starts in late elongating germ bands and is restricted to a domain within the neurogenic region throughout the depicted stages (**D_1_**_-**4**_). (**E_0_**_-**1**_) *Tc-ser* is partially co-expressed with *Tc-foxQ2* within a small sub-region of the AMR at elongating germ band stages (**E_0-1_**). (**E_2_**_-**4**_) At later stages the co-expression is restricted to outer lateral parts of the labral buds.

### *Tc-foxQ2* is required for *Tc-six3* expression and is repressed by Wnt signaling

We wondered in how far the interactions of *foxQ2* in other species were conserved in *Tribolium*. Unexpectedly, *Tc-six3* expression was strongly reduced or absent in *Tc-foxQ2*^RNAi^ germ rudiments (compare Fig. 7A_1_ with B_1_), which contrasts the findings in cnidarians and sea urchins. At later stages *Tc-six3* expression emerged but was reduced in strength and size with respect to the median expression domain (Fig. 7B_2-4_; empty arrowheads). The lateral neuroectodermal domains appeared to be more sensitive to *Tc-foxQ2* knockdown and were strongly reduced or even absent at later stages (Fig. 7B_2-4_; arrows). Conversely, *Tc-foxQ2* was virtually absent in *Tc-six3* RNAi embryos (Fig. 7D_1-4_) indicating a conserved role of *Tc-six3* in *Tc-foxQ2* regulation. At early embryonic stages, *Tc-wg* expression was not altered in *Tc-foxQ2*^RNAi^ (not shown). Likewise, *Tc-foxQ2* was not altered at early embryonic stages in *Tc-arr*^RNAi^ embryos, where ligand dependent canonical Wnt signaling is inhibited (Bolognesi et al., 2009). At later stages, however, the lateral neuroectodermal *Tc-foxQ2* domain was expanded in *Tc-arr*^RNAi^ (white arrows in Fig. 7E_2-4_) indicating a repressive function. In contrast, a mutual activation of *Tc-foxQ2* with *Tc-wg* was found at these later stages in the developing labrum (empty arrows in Fig. 7E_2-3_ and Fig. S6). In summary, in early embryos we found the expected activation of *Tc-foxQ2* by *Tc-six3* known from sea urchins and cnidarians. In contrast to these clades, however, *Tc-foxQ2* was required for *Tc-six3* expression. Based on *Tc-arr*^RNAi^ results, early *Tc-foxQ2* expression appears to be independent of ligand dependent Wnt signaling.

**Figure 7:**
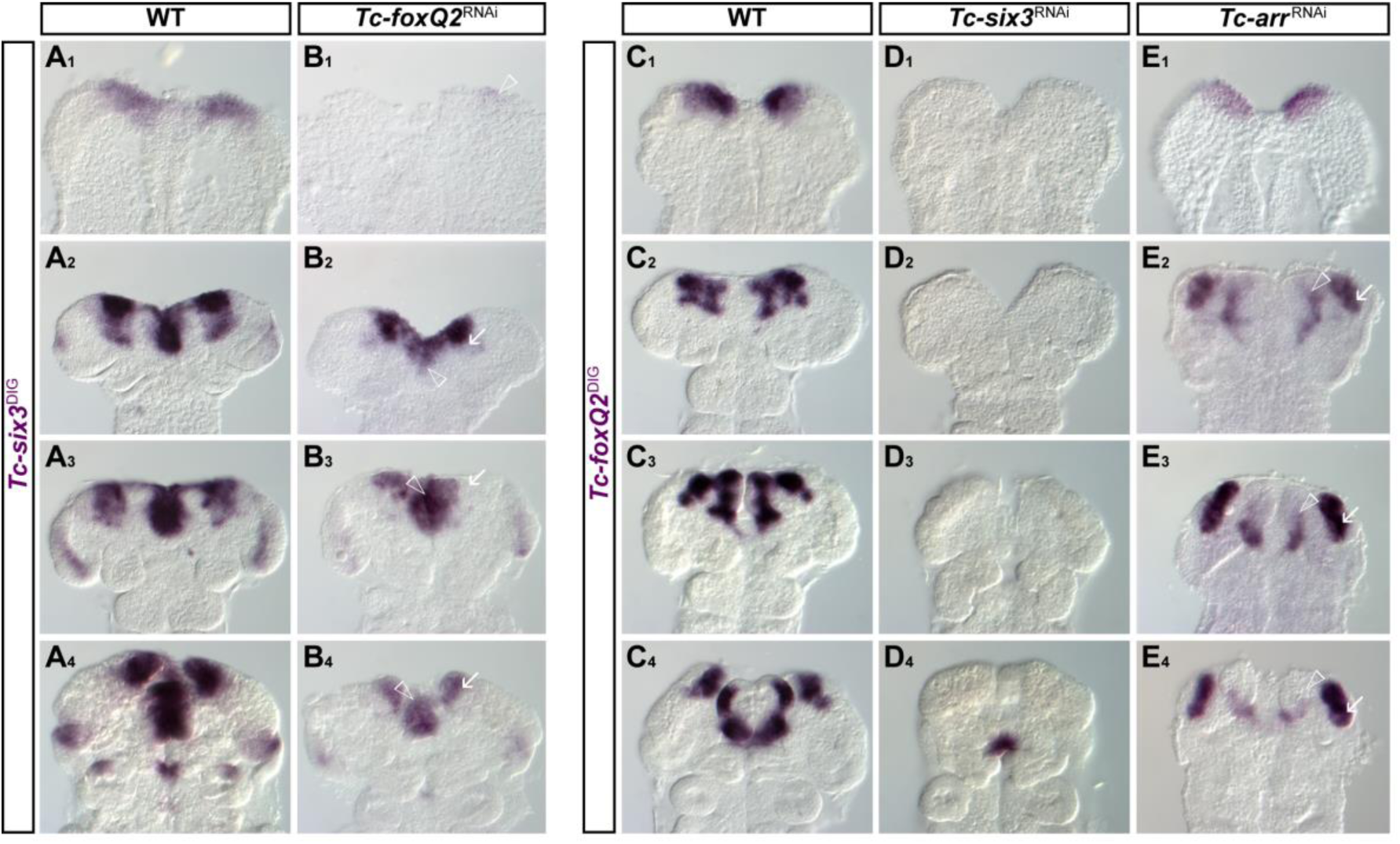
Mutual activation of *Tc-foxQ2* and *Tc-six3* and differential Wnt signaling-dependent *Tc-foxQ2* regulation. Anterior is up. Expression pattern of *Tc-six3* in wt (**A_1_**-**A_4_**) and *Tc-foxQ2*^RNAi^ (**B_1_**-**B_4_**) embryos and expression pattern of *Tc-foxQ2* in wt (**C_1_**-**C_4_**), *Tc-six3*^RNAi^ (**D_1_**-**D_4_**), and *Tc-arr*^RNAi^ (**E_1_**-**E_4_**) embryos monitored by ISH. (**B_1_**) *Tc-six3* expression is strongly reduced at *Tc-foxQ2*^RNAi^ germ rudiment stages (empty arrowhead). (**B_2_**) At later stages the median (empty arrowhead) and neurogenic (arrow) *Tc-six3* expression domains are reduced in size. (**B_3-4_**) At fully elongated germ band and early retracting germ band stages the labral (empty arrowhead) and the neuroectodermal (arrow) aspects of expression are strongly reduced, while the posterior median domain persists. The ocular domain appeared unchanged at late stages. (**D_1_**-**D_3_**) *Tc-foxQ2* expression is completely lost in early *Tc-six3*^RNAi^ embryos. (**D_4_**) The posterior portion of the stomodeal *Tc-foxQ2* expression domain emerges at retracting *Tc-six3*^RNAi^ germ band stages. (**E_1_**) *Tc-foxQ2* expression is unchanged at *Tc-arr*^RNAi^ germ rudiment stages. (**E_2_**-**E_4_**) At later stages the median *Tc-foxQ2* expression domain (empty arrowhead) is reduced, whereas the neurogenic *Tc-foxQ2* expression domain is expanded (arrow).

### *Tc-foxQ2* acts upstream in anterior AMR patterning

The anterior median region of the insect head (AMR) harbors the labrum and the stomodeum. *Tc-cnc* and *Tc-croc* are upstream factors required for anterior and posterior AMR patterning, respectively (Economou and Telford, 2009; Hunnekuhl and Akam, 2014; Kittelmann et al., 2013). The AMR expression domain of *Tc-cnc* in the labrum was strongly reduced after the knockdown of *Tc-foxQ2* (Fig. 8B_1-4_). Likewise, *Tc-croc* expression was affected but the reduction was restricted to the anterior boundary of expression (empty arrowheads in Fig. 8D_1-4_). Its posterior expression around the stomodeum was largely unchanged. Conversely, in *Tc-croc* and *Tc-cnc* RNAi we observed no alteration of *Tc-foxQ2* expression at early stages (not shown) indicating an upstream role of *Tc-foxQ2*. However, at later stages, expression of *Tc-foxQ2* was reduced in the labrum in both treatments (Fig. S7). Next, we tested the median AMR markers *Tc-scro*/*nk2.1* and *Tc-fkh*. *Tc-scro*/*nk2.1* was reduced anteriorly and laterally in *Tc-foxQ2*^RNAi^ embryos in early elongating germ bands (Fig. 8F_1_, empty arrowhead) but its posterior aspects remained unchanged. In contrast to wt embryos, the stomodeal/labral expression remained connected to the lateral expression in neuroectoderm (Fig. 8F_2-4_; empty arrow). Conversely, *Tc-foxQ2* was not changed in early *Tc-scro*/*nk2.1*^RNAi^ embryos (Fig. S7) while in later embryos, changes were observed. The expression of the stomodeum marker *Tc-fkh* was not considerably altered in *Tc-foxQ2*^RNAi^ embryos (Fig. S8) while later aspects of *Tc-foxQ2* expression were altered in *Tc-fkh* RNAi probably by indirect effects (Fig. S8).

**Figure 8:**
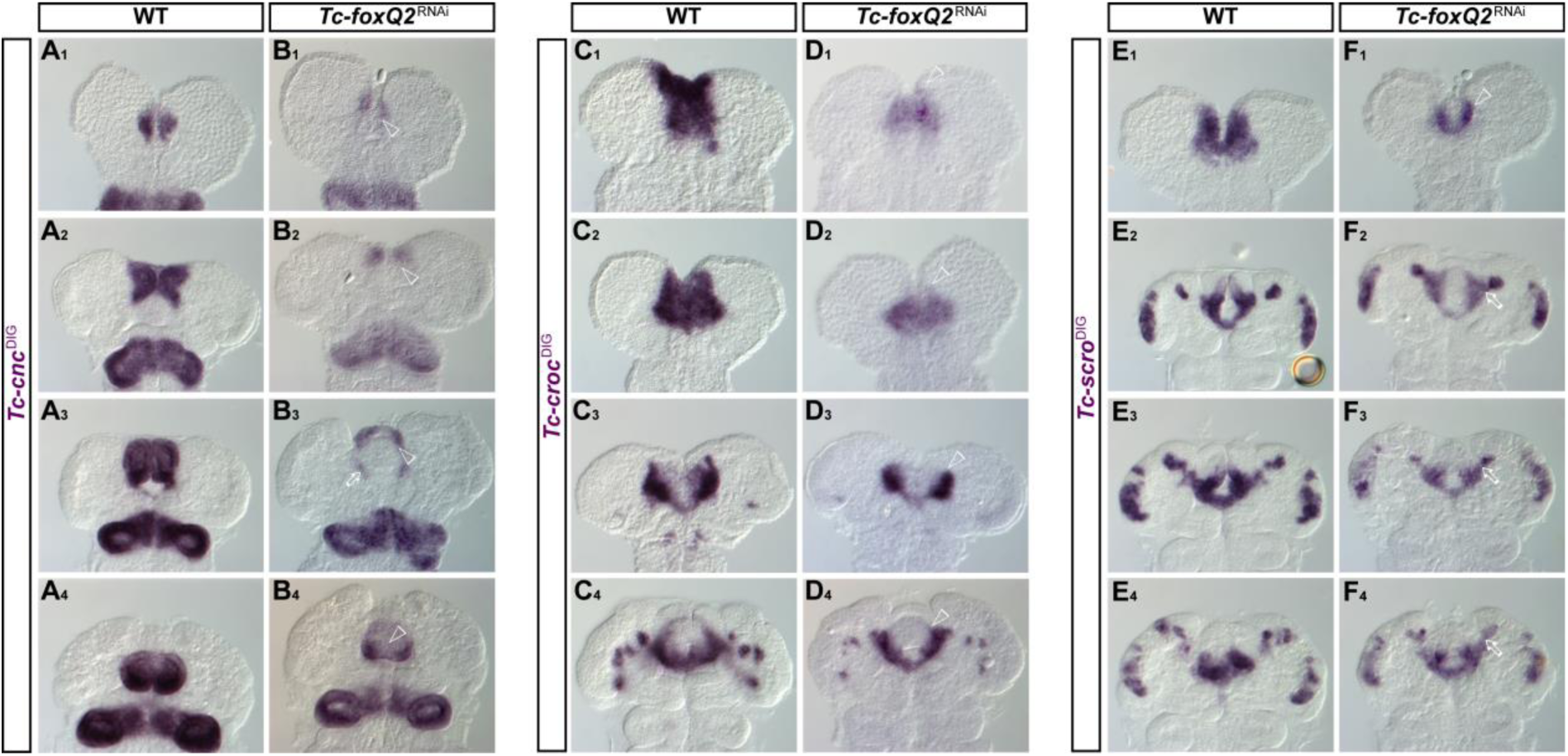
*Tc-foxQ2*^RNAi^ embryos show reduced *Tc-cnc* and *Tc-croc* expression domains and altered *Tc-scro* expression. Anterior is up. Expression pattern of *Tc-cnc* in wt (**A_1-4_**) and *Tc-foxQ2*^RNAi^ (**B_1_**_-**4**_) embryos, expression of *Tc-croc* in wt (**C_1_**_-**4**_) and *Tc-foxQ2*^RNAi^ (**D_1_**_-**4**_) embryos, as well as expression of *Tc-scro* in wt (**E_1_**_-**4**_) and *Tc-foxQ2*^RNAi^ (**F_1_**_-**4**_) embryos monitored by ISH. (**B_1_**_-**2**_) In *Tc-foxQ2*^RNAi^ embryos the AMR expression domain of *Tc-cnc* is reduced posteriorly during germ band elongation (empty arrowheads). Prior to this stage no considerable changes in the expression pattern were observed. (**B_3_**) Fully elongated germ bands show reduction in the labral buds whereas the stomodeal expression domain appears to be only slightly decreased (empty arrow). (**B_4_**) In retracting germ bands the expression of *Tc-cnc* in the anterior and median region of the labral buds is strongly reduced (empty arrowhead). (**D_1_**_-**4**_) Throughout development, *Tc-croc* expression pattern is lacking the anterior portion of its AMR domain (empty arrowheads). (**F_1_**) Expression of *Tc-scro*/*nk2.1* is reduced to a narrow stripe along the anterior fold (empty arrowhead). (**F_2_**_-**4**_) Later stages show an atypical bridging between the labral/stomodeal and the neurogenic *Tc-scro*/*nk2.1* expression domains (empty arrows).

The Notch pathway ligand *Tc-ser* and the ubiquitin ligase *Tc-mind bomb1* (*Tc-mib1*) are required for labrum development and knockdown of both lead to identical phenotypes (Siemanowski et al., 2015). However, we detected no difference in *Tc-ser* expression in *Tc-foxQ2*^RNAi^ (not shown). Conversely, in early and intermediate elongating *Tc-mib1*^RNAi^ embryos only the lateral aspects of *Tc-foxQ2* expression appeared mildly decreased (Fig. S9) arguing against a strong interaction with the Notch pathway. At later stages, in contrast, some lateral and labral *Tc-foxQ2* expression domains were clearly reduced in *Tc-mib1*^RNAi^ embryos. Taken together, these results demonstrated an upstream role of *Tc-foxQ2* in early anterior AMR patterning and they indicated that the later interactions of the aGRN are different from the early ones.

### An upstream role of *foxQ2* in neuroectoderm patterning

The anterior median neuroectoderm is marked by the expression of several highly conserved transcription factors and harbors the anlagen of the insect head placode (Posnien et al., 2011a), the pars intercerebralis and the pars lateralis (Posnien et al., 2011b) (Fig. S4). Further, it corresponds to the region where in grasshoppers several neuroblasts arise, which are required for CX development (Boyan and Reichert, 2011). *Tc-chx* expression was completely lost in early elongating *Tc-foxQ2*^RNAi^ germ bands (empty arrowhead in Fig. 9B_1_) and highly reduced at later stages (white arrows in Fig. 9B_2-4_). The *Tc-six4* domain was highly reduced in early *Tc-foxQ2*^RNAi^ embryos, showing only small spots of expression at the anterior rim (white arrow in Fig. 9D_1_). Later, the lateral expression developed normally while a median aspect of its expression was lost (white arrows in Fig. 9D_2-4_). *Tc-rx* expression at early elongating germ band stages was absent after *Tc-foxQ2*^RNAi^ (Fig. 9F_1_). This was unexpected because *Tc-rx* is largely expressed outside the *Tc-foxQ2* expression domain (Fig. 6A_0-2_) arguing against a direct effect. Indeed, our misexpression studies indicate a repressive role (see below). At later stages, the lateral aspects of *Tc-rx* expression recovered but the labral expression domain was reduced or lost, in line with co-expression (empty arrowheads in Fig. 9F_2-3_).

**Figure 9:**
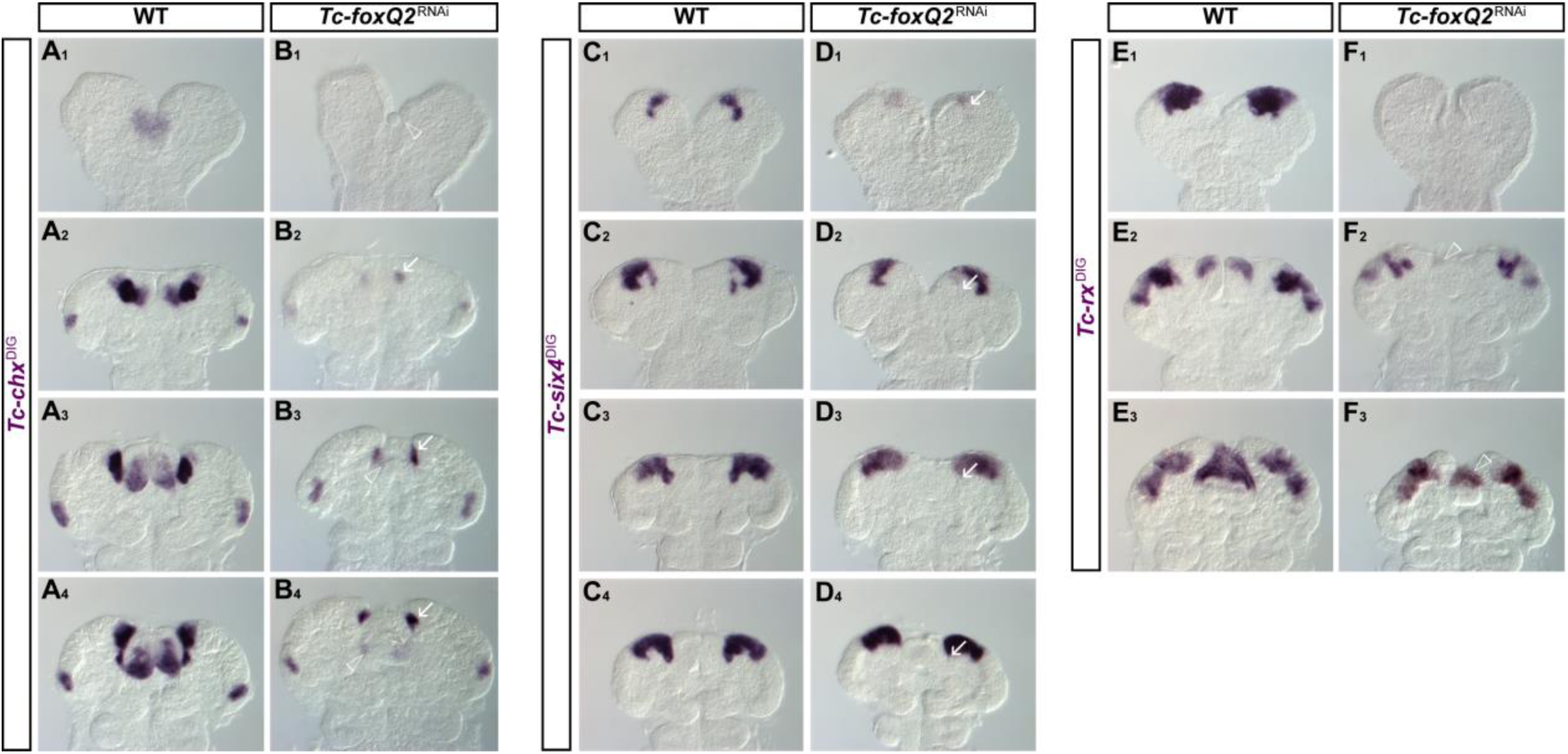
*Tc-foxQ2*^RNAi^ embryos show reduced *Tc-chx*, *Tc-six4*, and *Tc-rx* expression domains. Anterior is up. Expression patterns of *Tc-chx* in wt (**A_1_**_-**4**_) and *Tc-foxQ2*^RNAi^ embryos (**B_1_**_-**4**_), of *Tc-six4* in wt (**C_1_**_-**4**_) and *Tc-foxQ2*^RNAi^ embryos (**D_1_**_-**4**_) and of *Tc-rx* in wt (**E_1_**_-**3**_) and *Tc-foxQ2*^RNAi^ embryos (**F_1_**_-**3**_) monitored by ISH. (**B_1_**) *Tc-chx* expression is completely absent in early elongating *Tc-foxQ2*^RNAi^ germ bands (empty arrowhead). (**B_2_**_-**4**_) At later stages, the labral *Tc-chx* expression domains are almost absent (empty arrowheads) while the anterior neurogenic expression domains are strongly reduced (arrows). The ocular *Tc-chx* expression domains remain unaffected. (**D_1_**) Expression of *Tc-six4* is strongly reduced in early elongating germ bands (arrow). (**D_2_**_-**4**_) At later stages, the median posterior extensions of the *Tc-six4* expression domains are reduced (arrows). (**F_1_**) *Tc-rx* expression is strongly reduced or completely absent in early elongating *Tc-foxQ2*^RNAi^ germ bands. (**F_2_**_-**3**_) At later stages the neurogenic *Tc-rx* expression pattern appears unaffected, but the labral expression domains are absent (**F_2_**: empty arrowhead) or reduced in size (**F_3_**: empty arrowhead).

Next, we scored *Tc-foxQ2* expression in *Tc-chx*^RNAi^, *Tc-six4*^RNAi^ and *Tc-rx*^RNAi^ embryos. In neither treatment the early aspects of *Tc-foxQ2* expression were affected while at later stages, we found expression differences within the neurogenic region (Fig. S10). These results confirm the upstream role of *Tc-foxQ2* in early anterior patterning and they confirm that the interactions of the aGRN at later stages differ from the early ones. We found no change of cell death in the neurogenic region of *Tc-foxQ2*^RNAi^ embryos until retraction, where cell death was significantly increased 1.5 fold (*p*=0.023). Hence, the observed changes in expression domains are likely due to regulatory interactions but not due to loss of tissue (Fig. S11).

### *Tc-foxQ2* gain-of-function analysis confirms function in anterior median neuroectoderm

Heat shock-mediated misexpression has been established in *Tribolium* (Schinko et al., 2012) but has not yet been applied to the study of gene function. From eight independent transgenic lines we selected one for our experiments, which showed the most evenly distributed misexpression upon heat shock (Fig. S12). Heat shock misexpression led to a reproducible pleiotropic cuticle phenotype. With respect to the anterior head the bristle pattern showed diverse signs of mild disruption (Fig. S13). Outside its expression domain, *Tc-foxQ2* misexpression led to a reduced number of segments in the legs, the abdomen and the terminus. For our experiments, we used the earliest possible time point of misexpression (at 9-13 h AEL), which led to higher portion of anterior defects compared to 14-20 h AEL and 20-25 h AEL (not shown). At 14-18 h AEL the heat-shocked embryos were fixed and marker gene expression was scored. Wt embryos undergoing the same procedure were used as negative control.

Strongest effects were found with respect to genes with comparable late onset of expression, i.e. *Tc-rx*, *Tc-six4*, *Tc-scro*/nk2.1 and *Tc-cnc* (Fig. 10). The *Tc-rx* expression was reduced to a spotty pattern (Fig. 10A-C). This repressive function of *Tc-foxQ2* on *Tc-rx* is in line with the non-overlapping adjacent expression of these two genes (Fig. 6A_0-2_). Therefore, we assume that the reduction of *Tc-rx* found in RNAi is due to secondary effects rather (see above). Ectopic *Tc-foxQ2* expression caused a premature onset and an expansion of *Tc-six4* expression (Fig. 10E,F) and an additional ectopic domain was found in the posterior head (white arrowhead in Fig. 10E,F). *Tc-scro*/*nk2.1* expression emerged precociously and was expanded in size (Fig. 10H,I). Later, the domain was altered in shape and had a spotted appearance. In case of *Tc-cnc*, a posterior expansion of the expression was observed (empty arrowhead in Fig. 10N,O).

**Figure 10:**
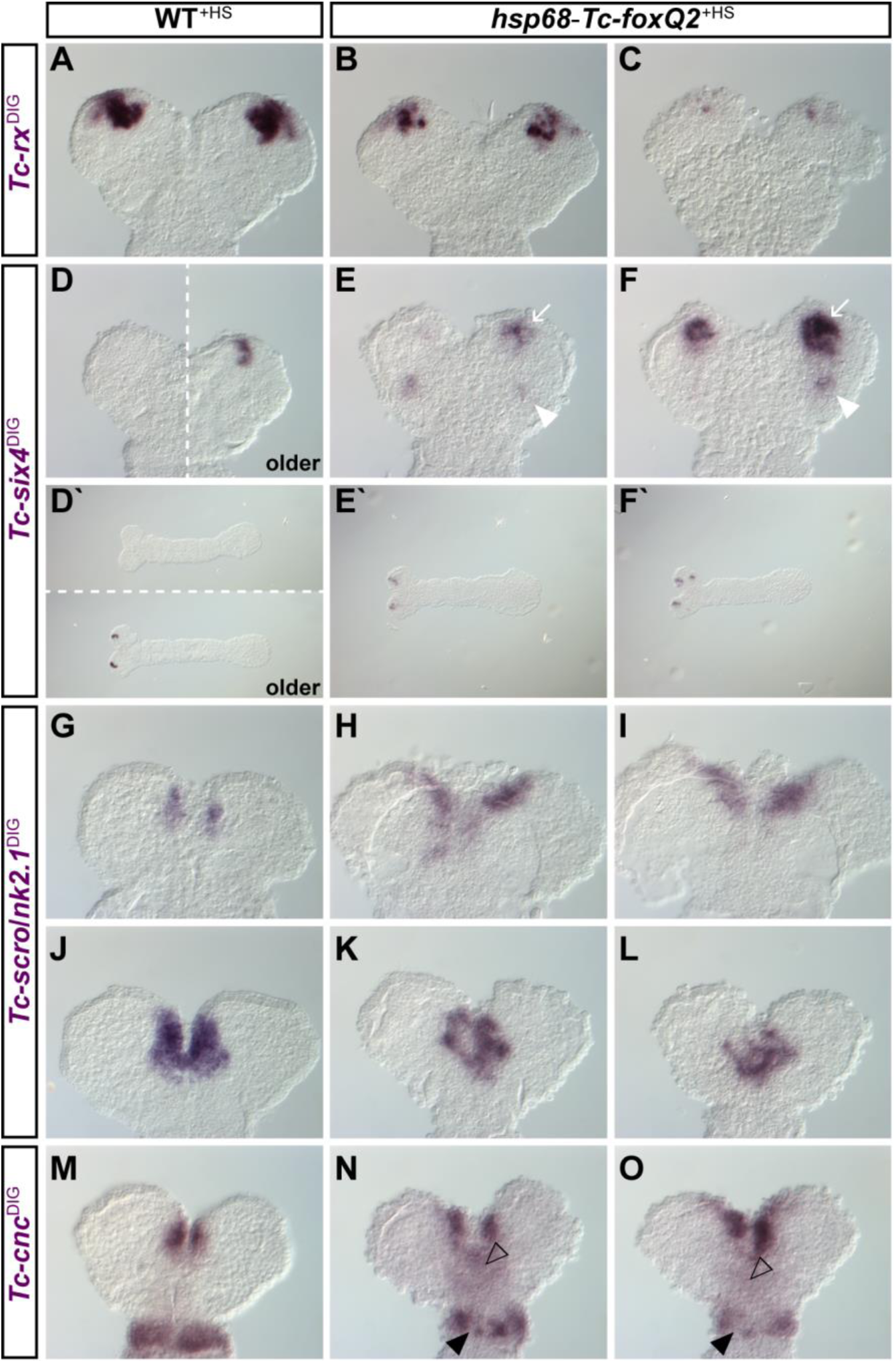
Ectopic *Tc-foxQ2* expression impacts head patterning gene expression profiles I. Anterior is up (left in **D**-**F**). Expression of head patterning genes in heat shock-treated wt (**A**, **D**, **D**, **G**, **J**, **M**) and *hsp68*-*Tc-foxQ2* (**B**, **C**, **E**, **E**, **F**, **F**, **H**, **I**, **K**, **L**, **N**, **O**) embryos (14-18 h AEL) is monitored by ISH. (**B**, **C**) Ectopic *Tc-foxQ2* expression leads to slightly (**B**) and heavily (**C**) reduced *Tc-rx* expression domains. (**E**, **E**, **F**, **F**) *Tc-six4* expression shows a premature onset (compare **E**, **F** with **D**) at the anterior tip (arrows), in *hsp68*-*Tc-foxQ2*^+HS^ embryos. These premature expression domains are expanded (**F:** arrow) compared to the size of the wt domains. Further, *hsp68*-*Tc-foxQ2*^+HS^ embryos show an additional *Tc-six4* expression domain within the antennal segment (white arrowhead). (**H**, **I**) The *Tc-scro*/*nk2.1* expression domains are, in *hsp68*-*Tc-foxQ2*^+HS^ germ rudiments, prematurely expressed and expanded. (**K**, **L**) In contrast, early elongating germ bands show reduced *Tc-scro*/*nk2.1* expression domains, in *hsp68*-*Tc-foxQ2*^+HS^ embryos. The later effect is presumably a secondary effect. (**N**, **O**) The anterior median *Tc-cnc* expression domains appear to be posteriorly spread (**N**: black empty arrowhead) and expanded in *hsp68*-*Tc-foxQ2*^+HS^ embryos (**O**: white empty arrowhead). The mandibular *Tc-cnc* expression domain is slightly (**N**) or heavily (**O**) reduced in a spotty manner, after ectopic *Tc-foxQ2* expression (black arrowheads).

Heat shock-induced expression starts only at late blastoderm stages (Schinko et al., 2012). Therefore, we were not able to test for the early interactions of the aGRN. Hence, comparably mild alterations of expression were found for *Tc-wg*, *Tc-six3* and *Tc-croc* after ectopic *Tc-foxQ2* expression (Fig. S14) in germ rudiments.

## Discussion

### Dynamic *six3/foxQ2* interactions govern anterior development

With this work we present the first functional analysis of *Tc-foxQ2* in any protostome and together with previous work we present the most comprehensive aGRN within protostomes (Fig. 11). We found that *Tc-foxQ2* is required at the top of the gene regulatory network to pattern the anterior-most part of the beetle embryo.

**Figure 11:**
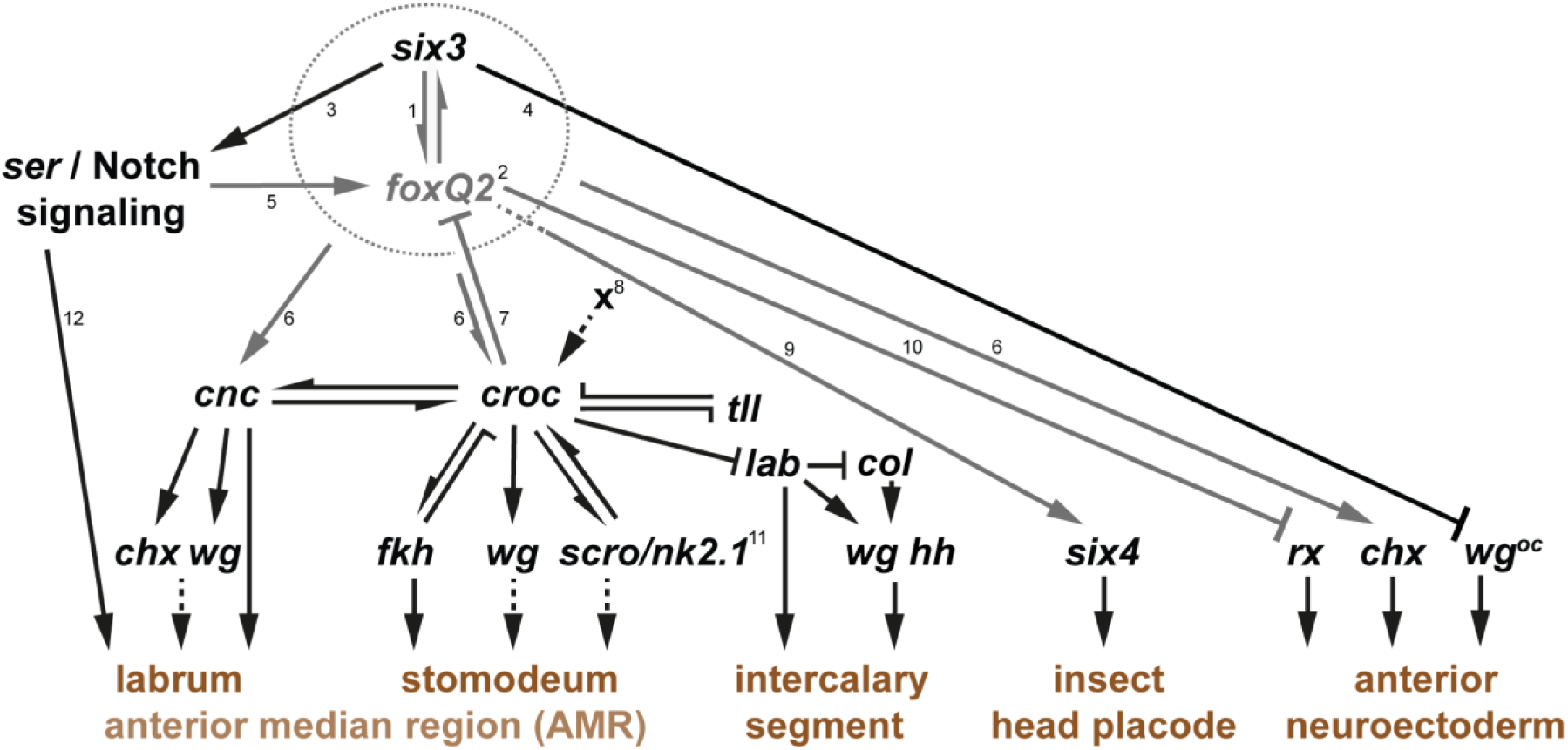
*Tc-foxQ2* forms an upstream regulatory module together with *Tc-six3*. Black lines indicate previously reported interactions (Based on (Kittelmann et al., 2013; Posnien et al., 2011c; Schaeper et al., 2010; Siemanowski et al., 2015)). Arrows represent gene activation, and cross-bars gene repression. Dashed lines indicate hypothetical effects. This aGRN represents the interactions between these genes at early embryonic stages – later interactions may differ (see text for details). (**1**) *Tc-six3* is the most upstream factor for patterning the anterior median head and neuroectoderm. (**2**) *Tc-foxQ2*, like *Tc-six3*, is a key player in anterior head development with a somewhat later onset of expression than *Tc-six3*, but similar cuticle phenotypes (epidermal & neural), and comparable activities in patterning of the anterior head. Mutual activation and similar phenotypes suggest that they form a regulatory module (indicated by the dashed circle). (**3**) *Tc-six3* acts on Notch signaling via *Tc-ser*. (**4**) *Tc-six3* prevents the ocular *Tc-wg* domain from expansion into the AMR, but is not acting on other *Tc-wg* domains. (**5**) Notch signaling-dependent activation of *Tc-foxQ2* is restricted to lateral parts of the anterior median *Tc-foxQ2* domains (*Tc-mib1* data). (**6**) Regulative activity is similar between *Tc-foxQ2* and *Tc-six3* with respect to several downstream targets. (**7**) *Tc-foxQ2* but not *Tc-six3* is repressed by *Tc-croc* activity. (**8**) An unknown factor ‘*X*’ is predicted to activate the posterior part of the *Tc-croc* expression, while *Tc-six3* and *Tc-foxQ2* are required for the anterior portion. (**9**) The interaction of *Tc-six3* with *Tc-six4* has not bee tested – hence it may or may not be regulated by the *Tc-foxQ2*/*Tc-six3* regulatory module (dashed line). (**10**) *Tc-rx* is repressed by *Tc-foxQ2* ectopic expression but is not regulated by *Tc-six3*. (**11**) The late effect of *Tc-foxQ2* on *Tc-scro*/*nk2.1*, observed in gain-of-function experiments, is most likely secondary and is, hence, not considered here. (**12**) Notch signaling is involved in labrum development by regulating cell proliferation.

Interestingly, we identified a positive regulatory loop between *Tc-six3* and *Tc-foxQ2* at early embryonic stages (germ rudiment) forming a novel regulatory module. In support of this, the epidermal and neural phenotypes of these genes are very similar as well as several genetic interactions (Posnien et al., 2011b). The *six3/foxQ2* regulatory module regulates a large number of genes that are required for anterior AMR patterning (e.g. anterior *Tc-cnc* and *Tc-croc* expression) and neuroectoderm patterning (e.g. *Tc-chx*). Hence, it lies at the top of the aGRN governing insect head and brain development. Such a self-regulatory module at the top of a gene network is not without precedence: For instance, *eyes absent*, *eyeless*, *dachshund,* and *sine oculis* form a positive regulatory loop at the top of the *Drosophila* eye gene regulatory network and as consequence all four genes are required for eye development leading to similar mutant phenotypes (i.e. complete loss of the eyes) (Wagner, 2007). With our data we are not able to distinguish between alternative modes of interactions: Both genes could cooperate in the regulation of the same targets, they could individually regulate different subsets, or one gene could be the regulator of all target genes while the other would just be required for initiation of expression.

Within the regulatory module, *Tc-six3* appears to be primus inter pares based on a number of criteria: Its expression starts a bit earlier (at the differentiated blastoderm stage compared to the germ rudiment expression of *Tc-foxQ2*) and has a larger expression domain, in which early *Tc-foxQ2* is completely nested. Further, the loss of *Tc-foxQ2* in *Tc-six3* RNAi is more complete even at later stages compared to the converse experiment (Fig. 7). Finally, the *Tc-six3* cuticle phenotype is more penetrant and comprises a slightly larger region of the dorsal head (compare with Posnien et al. 2011b).

Interestingly, the mutual activation of the *six3/foxQ2* module does not persist as the initially overlapping expression patterns diverge to become largely non-overlapping at later embryonic stages (Fig. 5B_2-6_). Just small domains in the neuroectoderm and the labrum continue to co-express both genes (encircled in Fig. 5B_3-6_). Given the almost mutually exclusive expression they might even switch to mutual repression at these later stages. In line with this scenario the heat shock misexpression of *Tc-foxQ2* led to a reduction of lateral aspects of *Tc-six3* expression (Fig. S14). Hence, it could be that later repression of *six3* by *foxQ2* is conserved between sea urchin and beetles.

### Protostome *foxQ2* evolved novel functions in head and brain development

Functional studies of *foxQ2* orthologs were restricted to a sea urchin as model for deuterostomes and the sea anemone, a cnidarian, representing the sister group to the bilaterian animals. We provide the first functional study in protostomes, which allows us drawing first conclusions on the evolution of *foxQ2* function. Compared to sea urchin and sea anemone, *Tc-foxQ2* plays a much more important role in our protostome model. First, it is clearly required for epidermal development documented in the loss of the entire labrum in knockdown animals. This is in contrast to sea urchin and sea anemone where no epidermal phenotype was described apart from a thickened animal plate (Sinigaglia et al., 2013; Yaguchi et al., 2008). Second, *Tc-foxQ2* is required for the development of two brain parts required for higher order processing, namely the CX and the MBs. Again, the neural phenotype in the other models was much weaker in that it affected the specification of certain cell types rather than the loss of tissues or brain parts (Sinigaglia et al., 2013; Yaguchi et al., 2008; Yaguchi et al., 2010). Finally, *Tc-foxQ2* is required for *Tc-six3* expression in *Tribolium*, which is not the case in the other models (Range and Wei, 2016; Sinigaglia et al., 2013; Yaguchi et al., 2008). Much of these novel functions may be explained by *Tc-foxQ2* gaining control over *Tc-six3* expression in our protostome model system.

### The evolutionary scenario: *foxQ2* gaining functions in animal evolution

Based on previous expression data, it has been suggested that *foxQ2* orthologs played a role in anterior development in all animals and that this involved interaction with *six3* orthologs (Fig. 12) (Fritzenwanker et al., 2014; Hunnekuhl and Akam, 2014; Marlow et al., 2014; Martín-Durán et al., 2015; Santagata et al., 2012; Sinigaglia et al., 2013; Tu et al., 2006; Wei et al., 2009). In line with a conserved function, at early stages, *foxQ2* shows co-expression with *six3* in deuterostomes and cnidarians and a nested expression within protostomes. In all cases, *foxQ2* arises within the *six3* domain (Fig. 12 left column). At later stages, in contrast, expression is more diverse. Some species retain nested or co-expression with *six3* while other species develop *six3* negative/*foxQ2* positive domains similar to what we found in the beetle. In other species, the anterior-most region is cleared from expression of both genes (Fig. 12 right column). This variation is not clearly linked to certain clades indicating that the regulatory interactions at later stages may have evolved independently. In addition to the novel functions described above, there are conserved functional aspects as well: Initial activation of *foxQ2* by *six3* is found in all three functional model species. Repression of *six3* by *foxQ2* was found in *Strongylocentrotus* and *Tribolium* (at later stages) but not *Nematostella* (Range and Wei, 2016; Sinigaglia et al., 2013).

**Figure 12:**
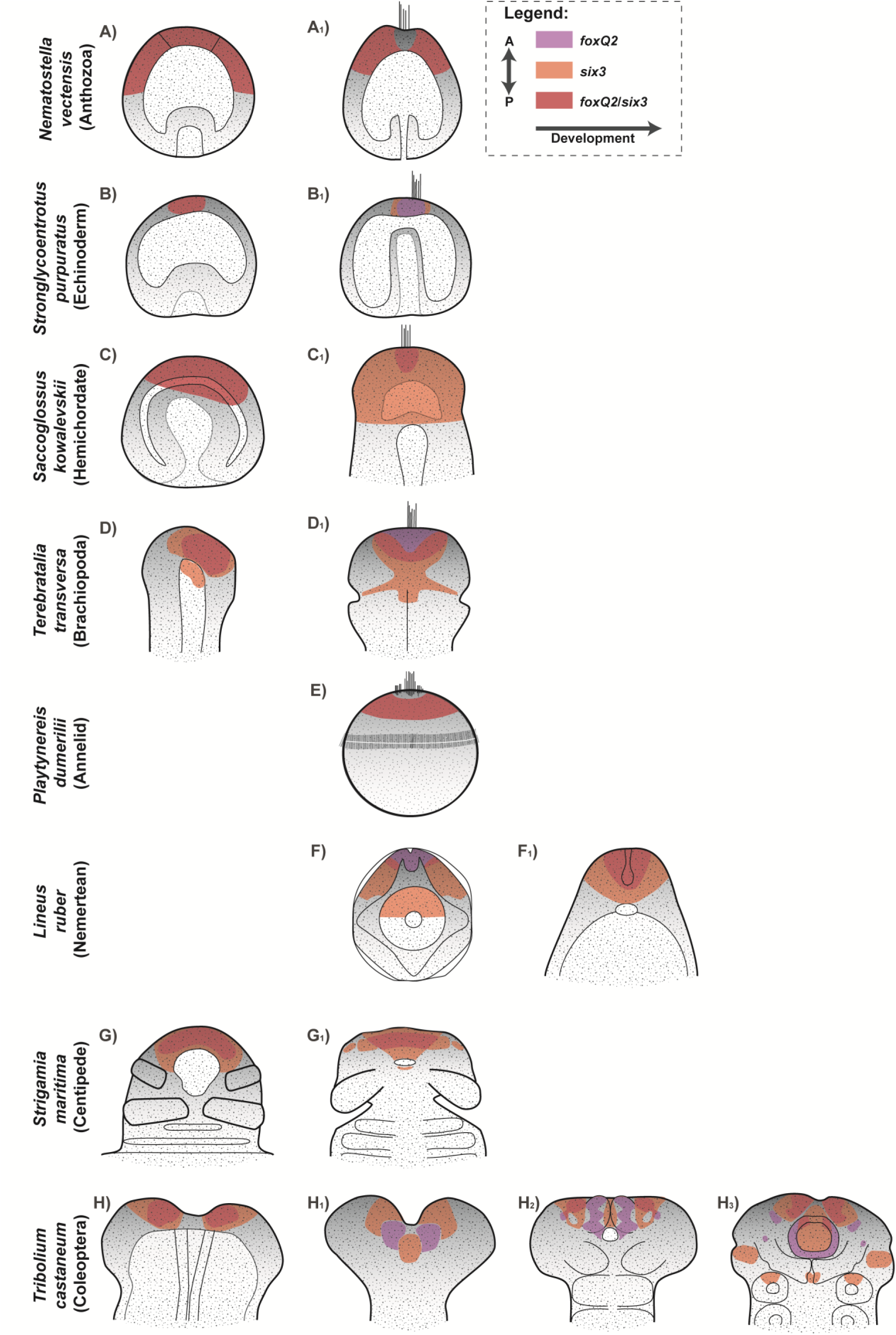
Expression of *foxQ2*/s*ix3* orthologs in different Metazoa. *foxQ2* (purple) and *six3* (orange) and their co-expression (red) at different developmental stages. The anterior/apical pole is oriented to the top. At early stages, co-expression of *foxQ2* and *six3* at the anterior pole of different metazoan species is highly conserved (left column). At later stages the patterns diverge leading to mutual exclusive expression in some taxa. (**A_1_**, **E**) *Nematostella* and *Platynereis* larvae show a *foxQ2*/*six3* coexpression during early stages like the other species, with the exception that the most apical region is free of *foxQ2*/*six3* expression. (**C_1_**, **F_1_**, **G_1_**) Late embryonic stages of *Saccoglossus* and *Strigamia* as well as early *Lineus* juveniles show a *foxQ2* expression at the anterior/apical pole, which is completely covered by *six3* expression. (**B_1_**, **H_1_**) *Strongylocentrotus* late gastrulae and *Tribolium* elongating germ bands show mutually exclusive expression of *foxQ2* and *six3* at later stages. (**D_1_**, **F_1_**) Early tri-lobed *Terebratalia* larvae and *Lineus* Schmidt’s larvae show *foxQ2* expression at the anterior/apical pole overlapping only posteriorly with *six3*. (**H_2_**_-**3**_) Fully elongated and retracting *Tribolium* germ bands show a complex expression pattern of *foxQ2* and *six3* with partial overlaps in the neuroectoderm (**H_2_**_-**3**_) and in the anterior labral buds (**H_3_**). (Based on (Fritzenwanker et al., 2014; Hunnekuhl and Akam, 2014; Marlow et al., 2014; Martín-Durán et al., 2015; Santagata et al., 2012; Sinigaglia et al., 2013; Tu et al., 2006; Wei et al., 2009); A: anterior, P: posterior

Together, these data indicate that at the base of metazoans *foxQ2* and *six3* were involved in early anterior patterning with *six3* being upstream of *foxQ2*. The anterior expression of both genes depended on repression by posterior Wnt signaling (Darras et al., 2011; Fritzenwanker et al., 2014; Marlow et al., 2014; Range and Wei, 2016; Sinigaglia et al., 2013; Wei et al., 2009; Yaguchi et al., 2008). After the split from Cnidaria, the Urbilateria *foxQ2* evolved a repressive function on *six3* leading to more complex expression and increased diversity of the molecular code specifying cells at the anterior pole (Range and Wei, 2016). This diversification may have been required for the evolution of more diverse neural cell types. In protostomes, *foxQ2* additionally evolved control over early *six3* expression. Indeed, the role of *six3* as primus inter pares in the regulatory module of *Tribolium* may be a remnant of the ancestrally more important role of *six3*. A curiosity is the loss of such a highly conserved gene in the genome of placental mammalians while other vertebrates still have the gene (Mazet et al., 2003; Yu et al., 2008). Unfortunately, *foxQ2* function in vertebrates remains unstudied. Of course, conclusions on gene function evolution based on one taxon for deuterostomes and protostomes, respectively, require further testing by analyses in other species. Hence, it will be interesting to see what the function of *foxQ2* is in other models representing deuterostomes, Lophotrochozoa and Ecdysozoa.

## Materials and Methods

### Animals and ethical statement

All the presented work is based on animals of the insect species *Tribolium castaneum*. Hence, an ethical approval was not required.

Animals were reared under standard conditions at 32°C (Brown et al., 2009). The *San Bernadino* (*SB*) wild-type strain was used for all experiments except for initial reproduction of the phenotype, where the *black* (Sokoloff, 1974) and the *Pig-19/pBA19* (Lorenzen et al., 2003) strains were used like in the iBeetle screen. The Tc-*vermillion*^*white*^(*v_w_*) strain (Lorenzen et al., 2002) was used for transgenesis and heat shock experiments. Transgenic lines marking parts of the brain (*MB-green* line (G11410); *brainy* line) were described in (Koniszewski et al., 2016).

### Sequence and phylogenetic analysis

*Tc-foxQ2* full coding sequence (1633 bp; Gen bank accession number: XM_008202469) was obtained from the *Tribolium* genome browser (http://bioinf.uni-greifswald.de/gb2/gbrowse/tcas5/) and the sequence was confirmed by cloning the full coding sequence from cDNA. Phylogenetic analysis was done by using MEGA v.5 (Tamura et al., 2011). The multiple sequence alignment was conducted with the ClustalW algorithm with the preset parameters. Positions containing gaps were eliminated from the dataset. The phylogenetic tree was constructed using the Neighbor-Joining method with the Dayhoff matrix based substitution model (Schwartz and Dayhoff, 1979). Bootstrap tests (Felsenstein, 1985) were conducted using 1000 replicates to test the robustness of the phylogenetic tree.

### RNAi

To test for RNAi efficiency, we detected *Tc-foxQ2* mRNA in *Tc-foxQ2* RNAi embryos (6-26 h AEL). As expected, no signal was detected using regular detection settings (not shown) but increasing the exposure time revealed residual *Tc-foxQ2* expression at advanced embryonic stages (Fig. S5; note the increased background in B,D,F,H). These domains reflected only part of the wt expression pattern indicating autoregulatory interactions restricted to some domains. Our subsequent analyses focused on early patterning where the RNAi knockdown was shown to be very efficient (Fig. S5A,B).

Template sequence of the dsRNA fragment *iB_03837* (Eupheria Biotech, Dresden, Germany) targeting *Tc-foxQ2,* which was used in the iBeetle screen:

5’-CAGCACATCCTCGACCACTATCCTTACTTCAGGACCCGGGGACCGGGTTGGAGAAACTCCATCAGGCATAATTTGTCTCTCAATGATTGTTTCATCAAGGCGGGAAGAAGCGCCAACGGAAAGGGACATTACTGGGCAATTCATCCCGCAAATGTGGACGACTTTAGAAAAGGGGACTTCAGGAGGAGGAAGGCACAAAGAAAGGTGAGGAAGCACATGGGGCTTGCCGTCGATGAAGATGGGGCTGATTCGCCAAGTCCGCCGCCCTTGTCTGTGAGTCCGCCTGTCGTGCCAGGGCCTTCCACGTCCGTTTATCACACAGTGCCGGCTCGAGGTCCGTCTCGCAAGCGGCAGTTCGACGTGGCGTCGCTTTTGGCGCCGGATTCCGGTGAAGACACCAACGAAGAGGACATCGACGTCGTCTCCAGTGACCAACACCAAGAGACTTCACCCAAACAGTGGCCTAATATGTTTCCCATCGTTAATTATTATCAAGCATTGTTACAAGCGA-3’.

The templates for the non-overlapping dsRNA fragments, used in this study, were generated by PCR from a plasmid template using following primers (including T7 RNA polymerase promoter sequence): Fragment *Tc-foxQ2*^RNAi_a^(489 bp): 5’-GAATTGTAATACGACTCACTATAGGCTTACTTCAGGACCCGG-3’ and 5’-GAATTGTAATACGACTCACTATAGGTCGCTTGTAACAATGCTTGA-3’; Fragment *Tc-foxQ2*^RNAi_b^, (197 bp): 5’-GAATTGTAATACGACTCACTATAGGATGTGCAGTAACGAGACTCC-3’ and 5’-GAATTGTAATACGACTCACTATAGGCTGGGGAAGAGCGGATAGC-3’. The dsRNA was synthesized using the Ambion^®^ T7 MEGAscript^®^ kit (lifeTechnologies, Carlsbad, CA, USA). The transcribed dsRNA was extracted via isopropanol precipitation (*Tc-foxQ2*^RNAi_a^) or phenol/chloroform extraction (*Tc-foxQ2*^RNAi_b^) and dissolved in injection buffer (1.4 mM NaCl, 0.07 mM Na_2_HPO_4_, 0.03 mM KH_2_PO_4_, 4 mM KCl, pH 6.8). The injected dsRNA concentrations for parental RNAi with *Tc-foxQ2*^RNAi_a^ and *Tc-foxQ2*^RNAi_b^ were 1.0 µg/µl, 1.5 µg/µl and 3.1 µg/µl. If not stated differently, a dsRNA concentration of 1.5 µg/µl was used. Pupal injections were performed as previously described (Bucher et al., 2002; Posnien et al., 2009b). The dsRNA was injected using FemtoJet^®^ express (eppendorf, Hamburg, Germany). Cuticles of the L1 larval offspring were prepared as described (Wohlfrom et al., 2006).

### Staining

Standard immunostaining was performed using the cleaved *Drosophila* Dcp-1 (Asp216) rabbit antibody (Cell Signaling Technology, Danvers, MA, USA; #9578) with 1:100 dilution. Anti-rabbit coupled with Alexa Fluor 488 was used for detection with 1:1000 dilution. ISH (alkaline phosphatase/NBT/BCIP) and DISH (alkaline phosphatase/NBT/BCIP & horseradish peroxidase mediated tyramide signal amplification (TSA) reaction) (Dylight550 conjugate synthesized by Georg Oberhofer) were performed as described previously (Oberhofer et al., 2014; Schinko et al., 2009; Siemanowski et al., 2015).

### Quantification of apoptosis

The regions of interest were set based on head morphology. Cell counting was performed using the Fiji cell counter plug-in (Schindelin et al., 2012). The number of apoptotic cells was positively correlated with age. Hence, to circumvent systematic errors due to staging the apoptotic cell number in the posterior procephalon was used to normalize the data. This region was chosen, because it was outside the *foxQ2* expression domain and unaffected by our RNAi experiments. The correction value was calculated by dividing the mean number of apoptotic cells of RNAi embryos by the mean number of apoptotic cells in wt embryos in the control region. For the normalization, the data was divided by the correction value. Raw counts are shown in Table S7.

The normalized data was tested with R (v.2.14.2; http://www.R-project.org/) for the homogeneity of the variances via the box plot, and for normal distribution, via the Shapiro-Wilk test. To test for significance, three statistical tests were conducted: Welch t-test, two sample t-test and the Wilcoxon rank-sum test. All three tests showed the same levels of significance. Stated *p*-values are based on the Wilcoxon rank-sum test results.

### Transgenesis and heat shock

The *foxQ2* heat shock construct was based on the constructs developed by Schinko et al., 2012 and germline transformation was performed as described previously (Berghammer et al., 1999; Schinko et al., 2012) using the injection buffer used for RNAi experiments and the mammalian codon-optimized hyperactive transposase (Yusa et al., 2011) flanked by the *Tc-hsp68* sequences (gift from Stefan Dippel). All animals for heat shock experiments were kept at 32°C. Heat shock was performed as described previously (Schinko et al., 2012) for ten minutes at 48°C (for cuticle preparations: 0-24 h AEL, 9-13 h AEL, 14-20 h AEL, 20-25 h AEL; for ISHs: 9-13 h AEL (fixated 5 h later)).

### Image documentation and processing

Cuticle preparations and L1 larval brains were imaged as described (Posnien et al., 2011b; Wohlfrom et al., 2006) using the LSM510 or Axioplan 2 (ZEISS, Jena, Germany) and processed using Amira (v.5.3.2; FEI) using ‘voltex’ projections. Stacks were visualized as average or maximum projections using Fiji (v. 1.49i; (Schindelin et al., 2012)). All images were level-adjusted and assembled in Photoshop CS (Adobe) and labelled using Illustrator CS5 (Adobe).

## Acknowledgements

We thank Claudia Hinners for excellent technical help and Stefan Dippel for the hyperactive helper construct. The phenotype was initially identified in the iBeetle project (DFG research unit FOR1234) and the work was supported by DFG grants BU1443/11, BU1443/8 and the Göttingen Graduate School for Neurosciences, Biophysics, and Molecular Biosciences (GGNB).

## List of Symbols and Abbreviatons

AEL: after egg laying
aGRN: anterior gene regulatory network
AMR: anterior median region
arr: arrow
CB: central body
cnc: cap’n’collar
col: collier
croc: crocodile
CX: central complex
Dcp1: cleaved *Drosophila* death caspase-1
fkh: forkhead
foxa: forkhead box a
foxq2: forkhead box q2
MB: mushroom body
mib1: mindbomb 1
nk2.1: nk2 homeobox 1 (thyroid transcription factor1
rx: retinal homeobox
scro: scarecrow
ser: serrate
six3: sine oculis homeobox homolog 3
six4: sine oculis homeobox homolog 4
tll: tailless
TSA: tyramide signal amplification
wg: wingless
wnt1: int1 (wingless-related1)

## Supporting Information Captions

Supplementary Figures

Given are 14 supplementary figures including legends supporting the data shown in the manuscript.

Supplementary Tables

Listed are raw counts of the quantification of apoptosis and the numbers visualized in the charts of Figures 1 and 2 and Fig. S2.

## Notes

The authors declare no competing interests.

